# Knocking out non-muscle myosin II in retinal ganglion cells promotes long-distance optic nerve regeneration

**DOI:** 10.1101/625707

**Authors:** Xue-Wei Wang, Shu-Guang Yang, Chi Zhang, Jin-Jin Ma, Yingchi Zhang, Bin-Bin Yang, Yi-Lan Weng, Guo-Li Ming, Anish R. Kosanam, Saijilafu, Feng-Quan Zhou

## Abstract

In addition to altered gene expression, pathological cytoskeletal dynamics in the axon are another key intrinsic barrier for axon regeneration in the central nervous system (CNS). Here we showed that knocking out myosin IIA/B in retinal ganglion cells alone either before or after optic nerve crush induced marked and sustained optic nerve regeneration. Combined Lin28 overexpression and myosin IIA/B knockout led to synergistic promoting effect and long-distance axon regeneration. Immunostaining, RNA-seq and western blot analyses revealed that myosin II deletion did not affect known axon regeneration signaling pathways or the expression of regeneration associated genes. Instead, it abolished the retraction bulb formation and significantly enhanced the axon extension efficiency. The study provided clear evidence that directly targeting neuronal cytoskeleton was sufficient to induce strong CNS axon regeneration, and combining gene expression in the soma and modified cytoskeletal dynamics in the axon was a promising approach for long-distance CNS axon regeneration.

## Introduction

Axon regeneration in the mammalian central nervous system (CNS) has been a long standing and highly challenging issue in the biomedical research field. The current consensus is that there are two major reasons that neurons in the mature mammalian CNS do not regenerate their axons after injury. One is the hostile environment created by inhibitors in the scar tissues and degenerating myelin, and the other is the diminished intrinsic neural regeneration ability of mature CNS neurons (*1, 2*). Therefore, the widely accepted view is that combination strategies that target both intrinsic growth ability and inhibitory environment are likely the best option for successful CNS axon regeneration and function recovery. Early studies (*3–6*) using peripheral nerve graft transplants have shown that some mature CNS neurons, such as spinal cord neurons and retinal ganglion cells (RGCs), could regenerate their axons into the permissive nerve grafts, indicating clearly that these neurons still retain limited intrinsic regeneration ability. However, to date, many studies targeting selected inhibitory molecules resulted in no or very modest CNS regeneration (*7, 8*). A likely reason is that there are multiple classes of inhibitory molecules, potentially including unidentified ones, which inhibit axon regeneration via distinct cellular and molecular mechanisms. Thus, targeting a few inhibitory signals while leaving the others intact may not result in a permissive environment similar to that in the peripheral nerve grafts.

In contrast, studies targeting the intrinsic axon growth ability have produced very promising results. In the optic nerve regeneration model, for example, Pten, Socs3, Klf4 loss of function, and Lin28 gain of function all achieved strong optic nerve regeneration (*9–13*). However, tissue clearing and 3D imaging studies have revealed that many of the regenerating RGC axons make U-turns in the optic nerve, at the optic chiasm, or make wrong guidance decisions after the chiasm (*14, 15*). In the corticospinal tract (CST) regeneration model, although modulation of the intrinsic regeneration ability substantially enhanced axon regeneration, most regenerating axons still cannot pass the lesion site, likely due to the effects of inhibitory molecules at the injury site, especially glial scar-based inhibitors. For instance, Pten deletion has been shown to induce by far the strongest promoting effect on CST axon regeneration (*16*). However, the most robust promoting effect can only be achieved in young mice (< 1 month). A recent study (*17*) showed that Pten deletion-induced regeneration of CST axons beyond the injury site was greatly diminished in aged mice. Specifically, in 12-18-month-old mice, Pten deletion led to little, if any, CST regeneration beyond the injury site. One likely reason for the diminished effect in older animals was the increased response to the inhibitory CNS environment. Thus, developing a successful approach for stimulating regeneration of injured CST remains a challenge, especially in older animals. Together, a new strategy is needed to enable neurons with increased intrinsic axon growth ability to grow axons in the inhibitory environment more efficiently with fewer U-turns and unnecessary branching, and can cross the inhibitory boundary more efficiently.

Neuronal cytoskeleton is not only the major machinery that drives axon growth (*18–20*), but also the converging targets of most, if not all, inhibitory signaling pathways (*18, 20*). In other words, by directly manipulating growth cone cytoskeletal motility it is possible to interfere with how the growth cones respond to multiple inhibitory signals, regardless if these signals are from different inhibitors or downstream pathways. Indeed, our previous study (*21*) showed that knocking down or pharmacologically inhibiting non-muscle myosin IIA/B could allow regenerating sensory axons to grow straight, make less branches, and completely ignore chondroitin sulfate proteoglycans (CSPGs) and myelin-based inhibitors. The effects were much stronger than that of Rho kinase inhibitor. Mechanistically, inhibition of myosin IIA/B resulted in loss of lamellipodia and actin arc, which led to significantly enhanced microtubule protrusion towards the leading edge of the growth cone. As a result, the axon growth rate over permissive substrate was greatly accelerated, and halted axon growth over inhibitory substrates was immediately restarted. Here we examined if knocking out non-muscle myosin IIA/B in RGCs could promote optic nerve regeneration in vivo. The results showed that deleting myosin IIA/B alone was sufficient to induce robust and sustained optic nerve regeneration. Moreover, combination of myosin IIA/B knockout and Lin28 overexpression, which enhances the intrinsic axon regeneration ability of RGCs (*12*), led to synergistic promoting effect and long-distance optic nerve regeneration. Importantly, the promoting effect was independent of well-known signaling mediators of optic nerve regeneration, such as increased mTOR activity and GSK3β inactivation (*2*). RNA-seq and western blot analyses comparing wild type and myosin IIA/B knockout neurons showed no significant difference in the expression of known regeneration associated genes, indicating local effects in the axons. In support, detailed analyses of growth cone morphologies and axon trajectories revealed that myosin II deletion almost abolished the formation of retraction bulbs, a hallmark of failed axon regeneration, and significantly enhanced the axon extension efficiency in the optic nerve. Furthermore, knocking out myosin IIA/B after optic nerve crush similarly enhanced optic nerve regeneration, indicating potential translational application. Collectively, our study demonstrated clearly that manipulation of neuronal cytoskeleton alone was sufficient to promote significant CNS axon regeneration in vivo. The study also provided strong evidence that combining enhanced intrinsic regeneration ability in the neuronal soma with local manipulation of axonal cytoskeleton was a promising approach to induce long-distance CNS axon regeneration in vivo.

## Results

### Double knockout of non-muscle myosin IIA/B in RGCs led to significant and sustained optic nerve regeneration in vivo

Non-muscle myosin II consists of two essential light chains, two regulatory light chains, and a myosin II heavy chain (MHC). There are three different isoforms of MHCs in mammalian cells, IIA, IIB, and IIC, encoded by *Myh 9*,*10*, and *14* genes, respectively. Myosin IIA and IIB with MHC IIA/B (myosin IIA/B) are the major isoforms in neurons (*21*). In nerve growth cones, both myosin IIA/B are localized near the transition zone, where microtubules and actin filaments interact (*21*). Our previous study (*21*) has shown that pharmacological inhibition or double knockdown of myosin IIA/B in developing or regenerating sensory neurons drastically promoted sensory axon growth over two major inhibitory substrates, myelin extracts or CSPGs. Here we tested if myosin IIA/B loss of function could also induce axon regeneration in RGCs after optic nerve injury. To knock out both myosin IIA/B in RGCs, we crossed *Myh9^f/f^* and *Myh10^f/f^* mice to generate *Myh9^f/f^: Myh10^f/f^* mice (hereafter *myosin IIA/B^f/f^*), and injected AAV2-Cre into the vitreous humors of these mice. Wild type mice injected with AAV2-Cre were used in the control group. To examine viral vector infection rate, immunostaining of Cre recombinase in whole-mount retina was performed two weeks after the injection. The results showed that the infection rate in RGCs was about 90% (Fig. S1A, C). Immunostaining of retinal sections also showed a nice colocalization of Cre staining with Tuj1-positive RGCs (Fig. S1B). We also injected AAV2-Cre into tdTomato reporter mice to examine the efficiency of Cre-mediated gene recombination. Strong expression of tdTomato in RGCs was observed two weeks after AAV2-Cre injection (Fig. S2A, B), indicating successful gene recombination. Lastly, we examined if myosin IIA/B were indeed deleted in RGCs after AAV2-Cre injection. By western blot analysis of the whole retina tissue, we found that the protein level of myosin IIA was markedly reduced (Fig. S3A). Immunostaining of retinal sections with anti-myosin IIB antibody showed significantly reduced level of myosin IIB in RGCs (Fig. S3B, C). Together, these results demonstrated clearly that myosin IIA/B were successfully deleted in RGCs.

To determine how myosin IIA/B double knockout (dKO) in RGCs affected optic nerve regeneration, we performed optic nerve crush (ONC) 2 weeks after the viral injection. We first assessed optic nerve regeneration 2 weeks after the ONC. The regenerating RGC axons were labeled with anterogradely transported cholera toxin subunit B (CTB) conjugated with Alexa Fluor 594, which was injected into the vitreous humor 2 days prior to tissue harvest (Fig. 1A). The fixed optic nerves were first tissue cleared to be transparent as previously described (*12*) and then axons were imaged with confocal microscopy. The results showed that very limited optic nerve regeneration occurred in wild type mice infected with AAV2-Cre. In contrast, there was greatly enhanced optic nerve regeneration in myosin IIA/B dKO mice (Fig. 1A, B). In the majority of optic nerves, regenerating axons reached 750 μm from the crush site. To further determine if myosin IIA/B dKO led to sustained promoting effect on optic nerve regeneration, we assessed optic nerve regeneration 4 weeks after the ONC. We found that at 4 weeks post ONC myosin IIA/B dKO significantly increased not only the lengths but also the number of regenerating axons, indicating continued axon regeneration along time. Specifically, most optic nerves had regenerating axons reaching 2000 μm from the crush site (Fig. 1B). In addition to counting axon numbers at different distances from the crush site, we also quantified the lengths of the top 5 longest regenerating axons in each condition. The result showed that in control condition the lengths of the top 5 axons remained unchanged at 2 and 4 weeks, whereas the top 5 axons of myosin IIA/B dKO RGCs continued to grow from 2 to 4 weeks (Fig. 1C). Both quantification results demonstrated that knocking out myosin IIA/B in RGCs led to sustained optic nerve regeneration at the same speed up to 4 weeks. When RGC survival rates were assessed, myosin IIA/B dKO showed no effect (Fig. 1D, E), indicating that it promoted optic nerve regeneration by enhancing axon extension rather than protecting RGCs from ONC-induced cell death. Collectively, these results demonstrated clearly that knocking out myosin IIA/B was sufficient to induce robust and sustained optic nerve regeneration without affecting cell survival.

**Fig. 1.**
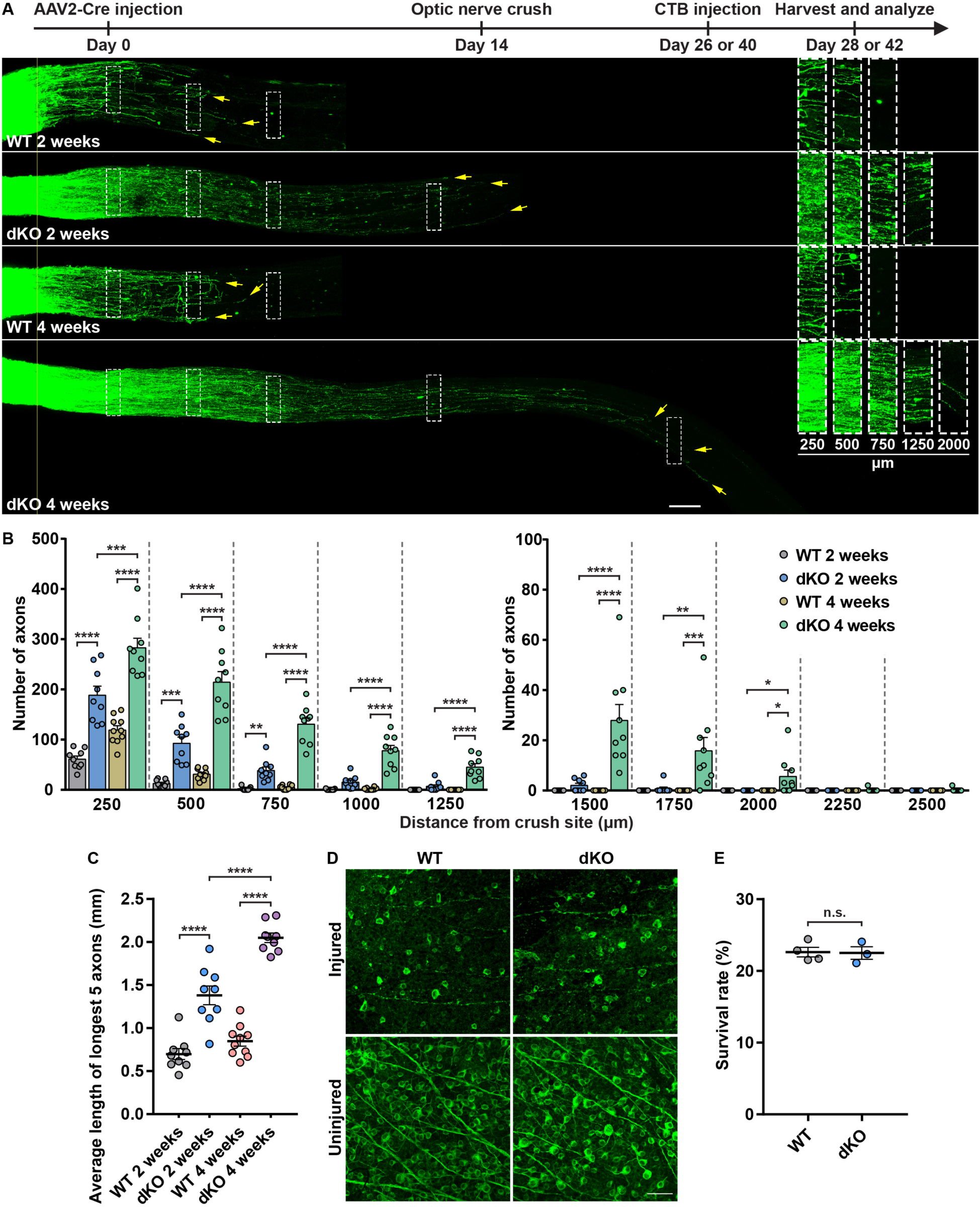
Deletion of myosin IIA/B in RGCs induced robust and sustained optic nerve regeneration. (A) Top: experimental timeline. Bottom: representative images of optic nerves showing that deletion of myosin IIA/B in RGCs produced robust and persistent axon regeneration 2 and 4 weeks after optic nerve crush. The columns on the right display magnified images of the areas in white dashed boxes on the left, showing axons at 250, 500, 750, 1250 and 2000 μm distal to the crush sites. The yellow line indicates the crush sites. Yellow arrows indicate the top 3 longest axons of each nerve. Scale bar, 100 μm (50 μm for the magnified images). (B) Quantification of optic nerve regeneration in (A) (one-way ANOVA followed by Tukey’s multiple comparisons test, *P* < 0.0001 at 250, 500, 750, 1000, 1250 and 1500 μm, *P* = 0.0002, 0.0071, 0.3875 and 0.3875 at 1750, 2000, 2250 and 2500 μm, respectively; n = 10 mice in 4-week WT group, n = 9 mice in other groups). (C) Quantification of the average length of the top 5 longest axons of each nerve in (A) (one-way ANOVA followed by Tukey’s multiple comparisons test, *P* < 0.0001; n = 10 mice in 4-week WT group, n = 9 mice in other groups). (D) Representative images of flat-mounted retinas showing that deletion of myosin IIA/B had no effect on RGC survival rate 2 weeks after optic nerve crush. Flat-mounted retinas were stained with anti-tubulin β3 antibody (Tuj1, green). Scale bar, 50 μm. (E) Quantification of RGC survival rate in (D) (unpaired t test, *P* = 0.9092, n = 4 and 3 mice in WT and dKO groups, respectively, 7-8 fields were analyzed for each retina). Data are represented as mean ± SEM. n.s., not significant, \**P* < 0.05, \*\**P* < 0.01, \*\*\**P* < 0.001, \*\*\*\**P* < 0.0001. WT, wild type; dKO, double knockout of myosin IIA/B.

### Knocking out non-muscle myosin IIA/B acted synergistically with Lin28a overexpression to promote long-distance optic nerve regeneration

Our recent study showed that overexpression of Lin28 in RGCs induced robust and sustained optic nerve regeneration via regulation of gene expression and enhanced intrinsic axon regeneration ability (*12*). Therefore, we tested if combining myosin IIA/B dKO, which alters axonal cytoskeletal dynamics, with Lin28 overexpression, which controls gene expression, could have combinatory promoting effect on optic nerve regeneration. The results showed that either myosin IIA/B dKO or Lin28a overexpression alone could lead to robust optic nerve regeneration 2 weeks after ONC (Fig. 2A, B). When both treatments were combined, optic nerve regeneration was greatly enhanced, with the longest length reaching 3.5mm from the crush site (Fig. 2A, B). In particular, either myosin IIA/B dKO or Lin28a overexpression RGCs had almost no regenerating axons growing beyond 1.75 mm from the crush site. In contrast, in the combinatory treatment group there were significant number of regenerating axons at 3 mm and the longest axons reached up to 3.5 mm (Fig. 2A, B). Because the dehydration process in the tissue clearing approach results in 18% of shrinkage in nerve lengths (*12*), the real lengths of the longest regenerating axons were more than 4 mm. Indeed, in about half of nerves in the combinatory group the longest regenerating axons almost reached the optic chiasm (Fig. 2A and Fig. S4) 2 weeks after ONC. This result suggested that myosin IIA/B dKO and Lin28a acted synergistically to promote long-distance optic nerve regeneration. Similarly, we quantified the average distances of the top 5 longest regenerating axons in each condition. The results showed that the longest regenerating axons in the combinatory group were markedly longer than either the myosin IIA/B dKO or Lin28a overexpression group (Fig. 2C). To better show the regenerating axons in the tissue cleared transparent optic nerves, we created a 3D animation of an optic nerve (Movie S1).

**Fig. 2.**
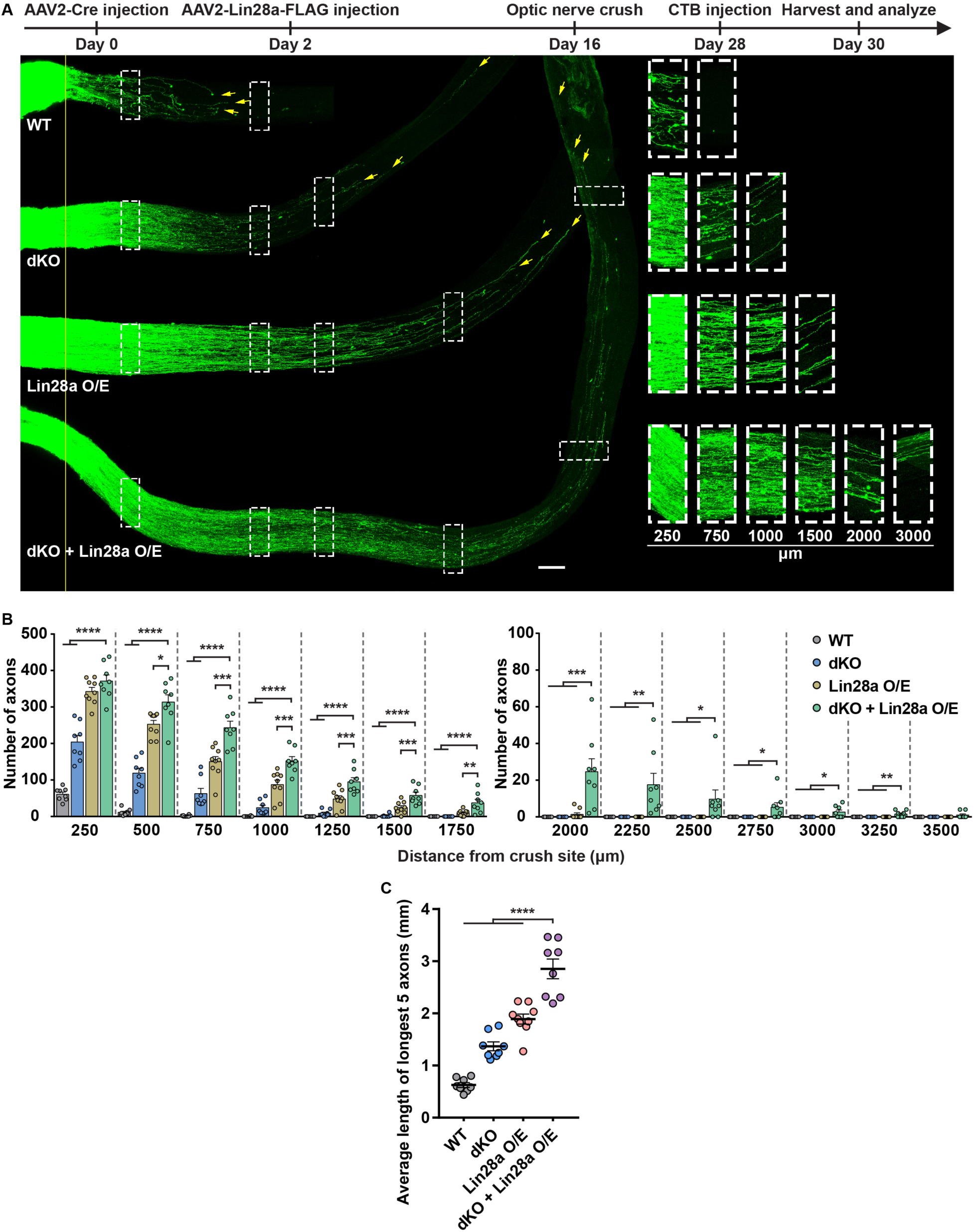
Myosin IIA/B deletion and Lin28a overexpression had synergistic effect on optic nerve regeneration. (A) Top: experimental timeline. Bottom: representative images of optic nerves showing that combining myosin IIA/B deletion with Lin28a overexpression in RGCs produced much stronger axon regeneration 2 weeks after the optic nerve crush. The columns on the right display magnified images of the areas in white dashed boxes on the left, showing axons at 250, 750, 1000, 1500, 2000 and 3000 μm distal to the crush sites. The yellow line indicates the crush sites. Yellow arrows indicate the top 3 longest axons of each nerve. Scale bar, 100 μm (50 μm for the magnified images). (B) Quantification of optic nerve regeneration in (A) (one-way ANOVA followed by Tukey’s multiple comparisons test, *P* < 0.0001 at 250, 500, 750, 1000, 1250, 1500, 1750 and 2000 μm, *P* = 0.0004, 0.0206, 0.0092, 0.0042, 0.0026, 0.0844 at 2250, 2500, 2750, 3000, 3250 and 3500 μm, respectively; n = 9 mice in Lin28a O/E group, n = 8 mice in other groups). (C) Quantification of the average length of the top 5 longest axons of each nerve in (A) (one-way ANOVA followed by Tukey’s multiple comparisons test, *P* < 0.0001; n = 9 mice in Lin28a O/E group, n = 8 mice in other groups). Data are represented as mean ± SEM. n.s., not significant, \**P* < 0.05, \*\**P* < 0.01, \*\*\**P* < 0.001, \*\*\*\**P* < 0.0001. WT, wild type; dKO, double knockout of myosin IIA/B; O/E, overexpression.

### Knocking out non-muscle myosin IIA/B in RGCs did not significantly affect known optic nerve regeneration signaling pathways

In our previous study (*21*), we showed that treating adult sensory neurons cultured on CSPGs with the myosin II inhibitor, blebbistatin, could induce halted axon to regrow within minutes. Conversely, washing out blebbistatin stopped axon growth in a very short time. Such rapid response to blebbistatin in both ways indicated that inhibition of myosin II promoted axon growth through its direct effects on the growth cone cytoskeleton without affecting signaling events in the neuronal soma. To test this idea, we examined how myosin IIA/B dKO affected two well-known pathways governing the intrinsic axon regeneration ability, the activation of mTOR, marked by increased level of phospho-S6 (pS6), and the inactivation of GSK3β, marked by phosphorylation of its serine 9 residue (pGSK3β), at 2 weeks after ONC. Previous studies have shown that most identified molecules promoting optic nerve regeneration act via either of these pathways, including Pten knockout (*9*), Akt overexpression (*22, 23*), Lin28 overexpression (*12*), osteopontin overexpression (*24*), SOCS3 deletion (*10*), melanopsin overexpression (*25*), HDAC5 manipulation (*26*), and direct modulation of mTOR (*27*) or GSK3β signaling (*22*). For mTOR activation, we examined the level of pS6 in RGCs under different conditions. The results showed that knocking out myosin IIA/B had no effect on pS6 level, whereas Lin28a overexpression markedly increased the level of pS6 in RGCs (Fig. 3A). Quantification demonstrated that the percentage of pS6 positive (pS6^+^) RGCs increased by nearly 8 folds in the Lin28a overexpression group compared to that in the wild type group, whereas myosin IIA/B dKO had no effect (Fig. 3B). To provide a more objective measurement of pS6 level in RGCs, we also quantified the average fluorescence intensity of pS6 staining in all Tuj1 positive RGCs. The results showed that myosin IIA/B dKO actually slightly reduced the pS6 level compared to that of the wild type group. In contrast, Lin28a overexpression greatly increased the pS6 level (Fig. 3C). Lastly, we quantified the average fluorescence intensity of pS6 staining only in pS6^+^ RGCs under different conditions. Similarly, there was no significant difference between wild type and myosin IIA/B dKO RGCs, whereas the Lin28a overexpression group had a much higher value (Fig. 3D).

**Fig. 3.**
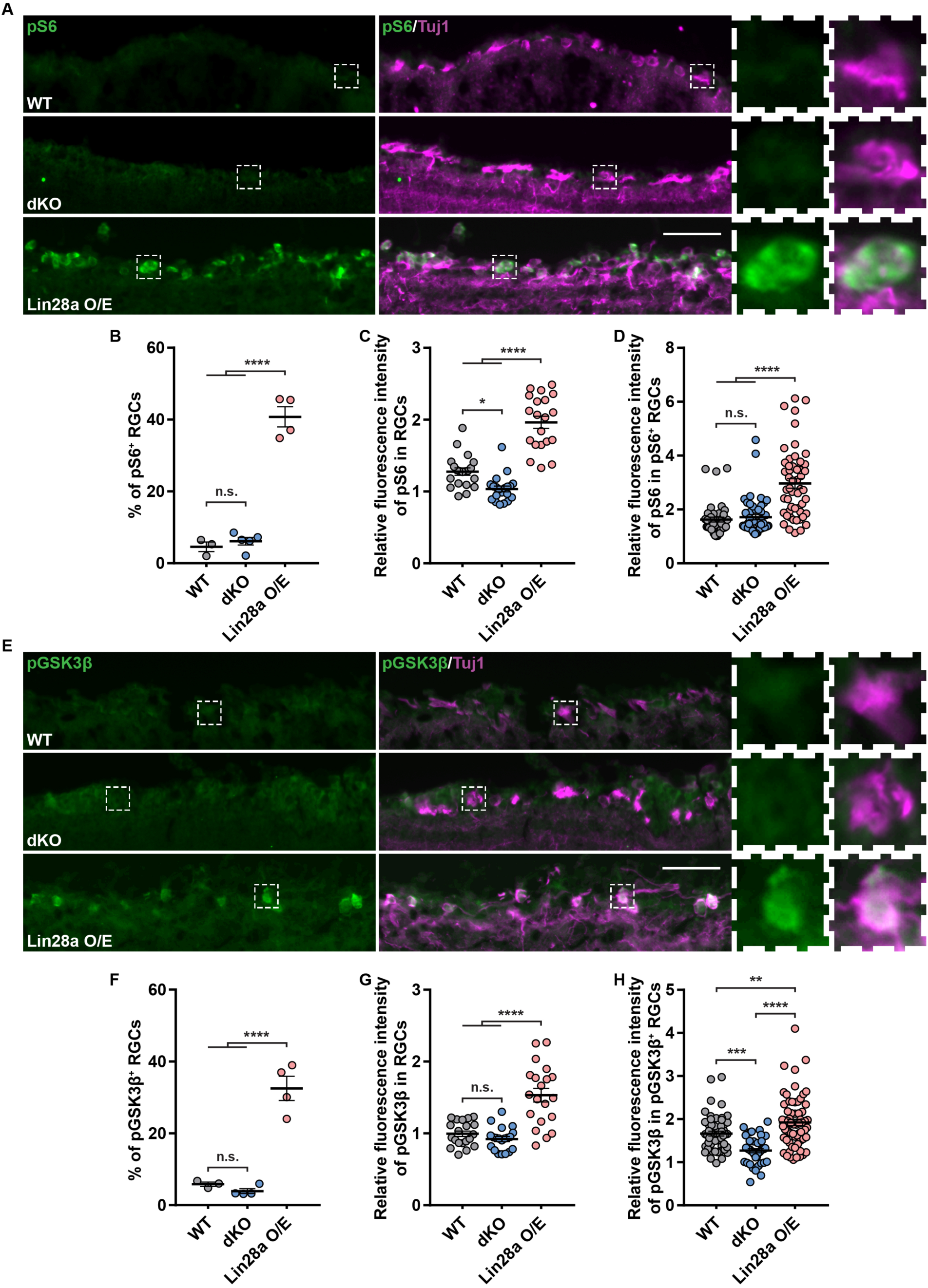
Myosin IIA/B deletion-induced optic nerve regeneration was independent of mTOR or GSK3β pathway. (A) Representative images of retinal sections showing that the deletion of myosin IIA/B did not activate mTOR (marked by pS6) in RGCs, whereas Lin28a overexpression markedly activated mTOR in RGCs 2 weeks after optic nerve crush. The two columns on the right display magnified images of the RGCs marked in white dashed boxes on the left. Retinal sections were stained with anti-pS6 (green) and anti-tubulin β3 (magenta) antibodies. Scale bar, 50 μm (12.5 μm for the magnified images). (B) Quantification of the percentage of pS6^+^ RGCs in (A) (one-way ANOVA followed by Tukey’s multiple comparisons, *P* < 0.0001, n = 3 mice in WT group, n = 5 mice in dKO group, n = 4 mice in Lin28a O/E group, at least 363 RGCs from at least 7 non-adjacent retinal sections were analyzed for each mouse). (C) Quantification of average fluorescence intensity of pS6 in all RGCs (one-way ANOVA followed by Tukey’s multiple comparisons, *P* < 0.0001, n = 20 retinal sections with identical imaging configurations from at least 2 mice were analyzed for each group). (D) Quantification of average fluorescence intensity of pS6 in pS6^+^ RGCs (one-way ANOVA followed by Tukey’s multiple comparisons, *P* < 0.0001, n = 42, 49 and 53 RGCs with identical imaging configurations from at least 2 mice were analyzed for WT, dKO and Lin28a O/E groups, respectively). (E) Representative images of retinal sections showing that the deletion of myosin IIA/B did not inactivate GSK3β (marked by pGSK3β) in RGCs, whereas Lin28a overexpression markedly inactivated GSK3β in RGCs 2 weeks after optic nerve crush. The right two columns display magnified images of the RGCs marked in white dashed boxes on the left. Retinal sections were stained with anti-pGSK3β (green) and anti-tubulin β3 (magenta) antibodies. Scale bar, 50 μm (12.5 μm for the magnified images). (F) Quantification of the percentage of pGSK3β^+^ RGCs in (E) (one-way ANOVA followed by Tukey’s multiple comparisons, *P* < 0.0001, n = 3 mice in WT group, n = 4 mice in other groups, at least 434 RGCs from at least 7 non-adjacent retinal sections were analyzed for each mouse). (G) Quantification of average fluorescence intensity of pGSK3β in all RGCs (one-way ANOVA followed by Tukey’s multiple comparisons, *P* < 0.0001, n = 20 retinal sections with identical imaging configurations from at least 2 mice were analyzed for each group). (H) Quantification of average fluorescence intensity of pGSK3β in pGSK3β^+^ RGCs (one-way ANOVA followed by Tukey’s multiple comparisons, *P* < 0.0001, n = 54, 40 and 65 RGCs with identical imaging configurations from at least 2 mice were analyzed for WT, dKO and Lin28a O/E groups, respectively). Data are represented as mean ± SEM. n.s., not significant, \**P* < 0.05, \*\**P* < 0.01, \*\*\**P* < 0.001, \*\*\*\**P* < 0.0001. WT, wild type; dKO, double knockout of myosin IIA/B; O/E, overexpression.

For GSK3β inactivation, we found that there were very few wild type RGCs showing positive staining of pGSK3β, and knocking out myosin IIA/B had no impact on it (Fig. 3E, F). In contrast, Lin28a overexpression increased the percentage of pGSK3β positive (pGSK3β^+^) RGCs by about 6 folds (Fig. 3F). The average fluorescence intensity of pGSK3β staining in all Tuj1 positive RGCs was not affected by myosin IIA/B dKO, whereas Lin28a overexpression greatly increased the pGSK3β level (Fig. 3G). When the average fluorescence intensity of pGSK3β was quantified only in pGSK3β^+^ RGCs, the level of pGSK3β was significantly decreased in myosin IIA/B dKO group compared with that in the wild type group. The level of pGSK3β in the Lin28a overexpression group was still the highest (Fig. 3H).

Taken together, these results provided clear and strong evidence that knocking out myosin IIA/B had no effects on two well-known signaling pathways occurred in the neuronal soma supporting intrinsic regenerative ability of optic nerves.

### Knocking out non-muscle myosin IIA/B in sensory neurons did not significantly affect known regeneration associated genes and pathways

Double knockdown of myosin IIA/B in sensory neurons drastically promoted regenerative axon growth over CSPGs or CNS myelin (*21*), similar to our finding in RGC axons. Therefore, we generated *Advillin-Cre: myosin IIA/B^f/f^* mice, in which myosin IIA/B were conditionally knocked out in sensory neurons, to explore if myosin IIA/B dKO in neurons affected known regeneration associated genes and pathways. We performed sham surgery or bilateral sciatic nerve injury (SNI) on these mice or wild type mice. Three days later, we collected lumbar 4 and 5 dorsal root ganglia (L4/5 DRGs) and isolated mRNA and protein for RNA-seq and western blot analyses. As expected, mRNA levels of myosin IIA/B in L4/5 DRGs of *Advillin-Cre: myosin IIA/B^f/f^* mice were lowered (Fig. 4A). Pearson correlation coefficient of gene expression indicated by FPKM (Fragments Per Kilobase of exon model per Million mapped reads) among different samples showed that under the same treatment (sham or SNI), the two wild type replicates were quite similar to the two dKO replicates (Fig. 4B). When the numbers of differentially expressed genes (DEGs) were compared, we found SNI significantly altered the mRNA levels of nearly 2000 genes, whereas myosin IIA/B dKO caused little change. Specifically, the number of DEGs between wild type neurons and myosin IIA/B dKO neurons was actually equivalent to that between the two replicates within each condition (Fig. 4C). These results indicated that myosin IIA/B dKO *per se* did not significantly change the transcriptome in sensory neurons. In addition, in wild type and myosin IIA/B dKO neurons, we closely examined and compared the FPKMs of many classic regeneration-associated genes (RAGs) and genes well-known to control axon regeneration, such as *Atf3*, *Sox11*, *Lin28a*, *Gap43*, *Pten*, *Klf9*, and *Rab27*, etc (*12, 28–32*). The results showed that the mRNA levels of these genes were up- or downregulated by SNI as expected, but were not largely affected by myosin IIA/B dKO (Fig. 4D, E). Gene ontology analysis of DEGs between different conditions revealed that SNI stimulated many neuron-related programs (Fig. 4G), whereas myosin IIA/B dKO induced gene expression change had no specific connection with neurons or axon regeneration (Fig. 4H). Lastly, we directly examined the protein levels of several genes and pathways regulating axon regeneration by western blot and found that in either uninjured or injured condition, myosin IIA/B dKO did not change the protein levels of Atf3, c-Jun, Gap43 or c-Myc (Fig. 4F). Moreover, consistent with our RGC immunostaining results, myosin IIA/B dKO had no impact on mTOR or GSK3β pathway (Fig. 4F) in sensory neurons. Taken together, these results further supported that myosin IIA/B dKO induced optic nerve regeneration was unlikely to be caused by enhanced intrinsic axon regeneration ability.

**Fig. 4.**
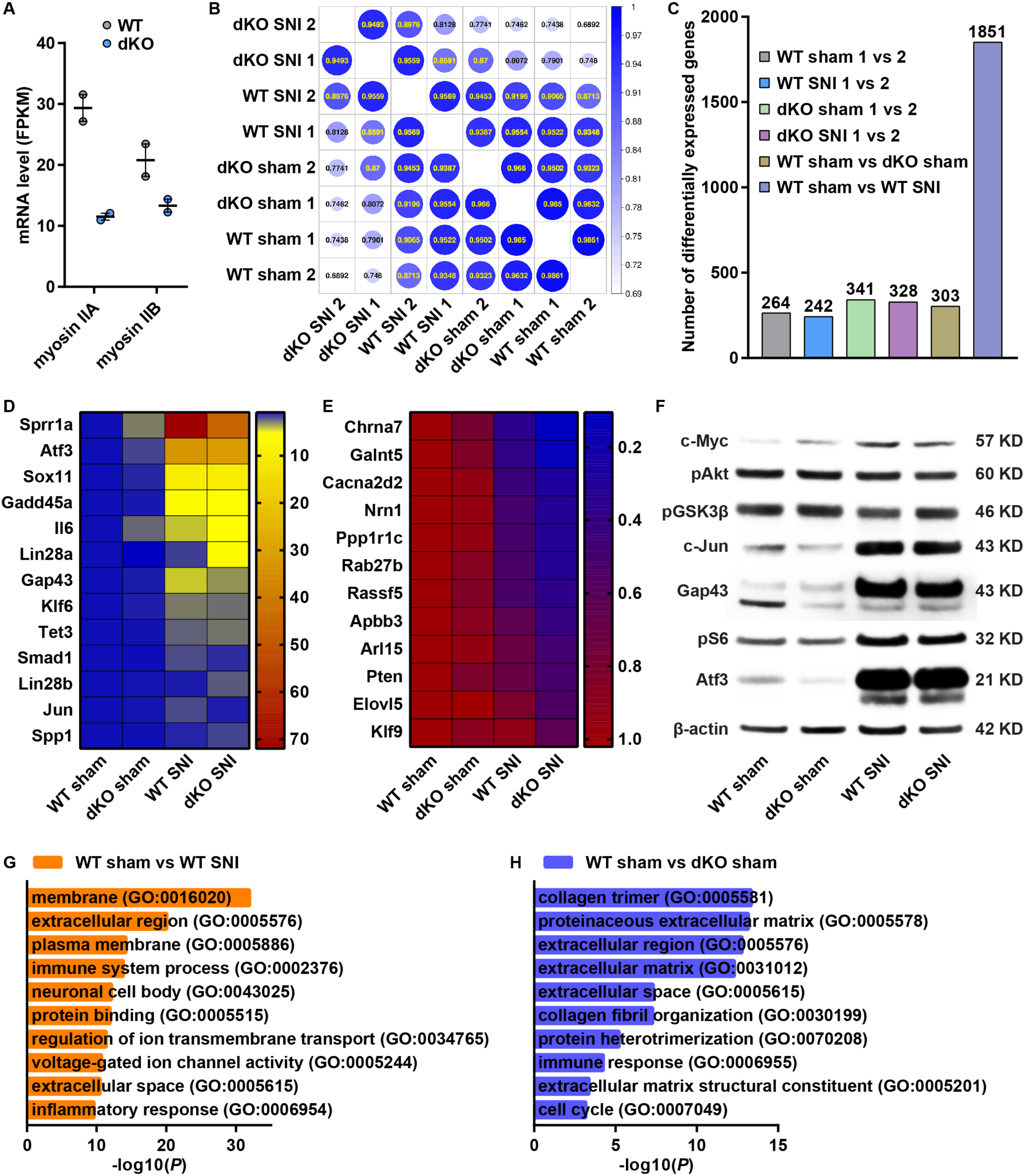
Conditional knockout of myosin IIA/B in sensory neurons did not significantly affect known regeneration associated genes and pathways. (A) Decreased mRNA levels of myosin IIA/B in L4/5 DRGs of *Advillin-Cre: myosin IIA/B^f/f^* mice (n = 2 mice in each group, data are represented as mean ± SEM). (B) Pearson correlation heatmap showing the high degree of similarity in gene transcription between wild type and myosin IIA/B deleted neurons under the same condition (sham or SNI). (C) Number of differentially expressed genes between replicates or conditions showing that myosin IIA/B deletion did not significantly change gene expression in neurons. (D, E) Expression heatmap of genes known to be upregulated (D) or downregulated (E) following SNI. Results showed that myosin IIA/B deletion did not affect the transcription of these genes in neurons. (F) Representative western blot results showing that neuron specific myosin IIA/B conditional knockout did not affect classic regeneration associated genes or pathways (n = 3 independent experiments). (G, H) Gene ontology analyses of differentially expressed genes between uninjured (sham) and injured (SNI) condition in wild type neurons (G), and between wild type neurons and myosin IIA/B deleted neurons under uninjured (sham) condition (H). WT, wild type. dKO, double knockout of myosin IIA/B. SNI, sciatic nerve injury.

### Knocking out non-muscle myosin IIA/B in RGCs changed axon tip morphology and regenerating axon trajectory

To better understand the cellular mechanisms by which myosin IIA/B knockout promoted optic nerve regeneration locally at the axons, we first performed detailed analysis of axonal tip morphologies in wild type and myosin IIA/B dKO optic nerves 2 and 4 weeks after ONC. Based on a previous study (*33*), there are mainly three types of dynamic axonal tip morphologies in vivo. One is the retraction bulb (Fig. 5A, C), which is the hallmark structure of dystrophic axons that failed to regenerate (*18, 20*). The other two are growth-competent growth cones with two different end shapes (Fig. 5A, C). We found that in wild type optic nerves, a significant percentage of axons had retraction bulbs at their ends, whereas knocking out myosin IIA/B in RGCs almost abolished the formation of retraction bulbs (Fig. 5A, B). Most regenerating axons in the myosin IIA/B knockout nerves had growth-competent growth cones, indicating that deleting myosin IIA/B efficiently transformed dystrophic axon tips into growth cones and rendered subsequent axon regeneration.

**Fig. 5.**
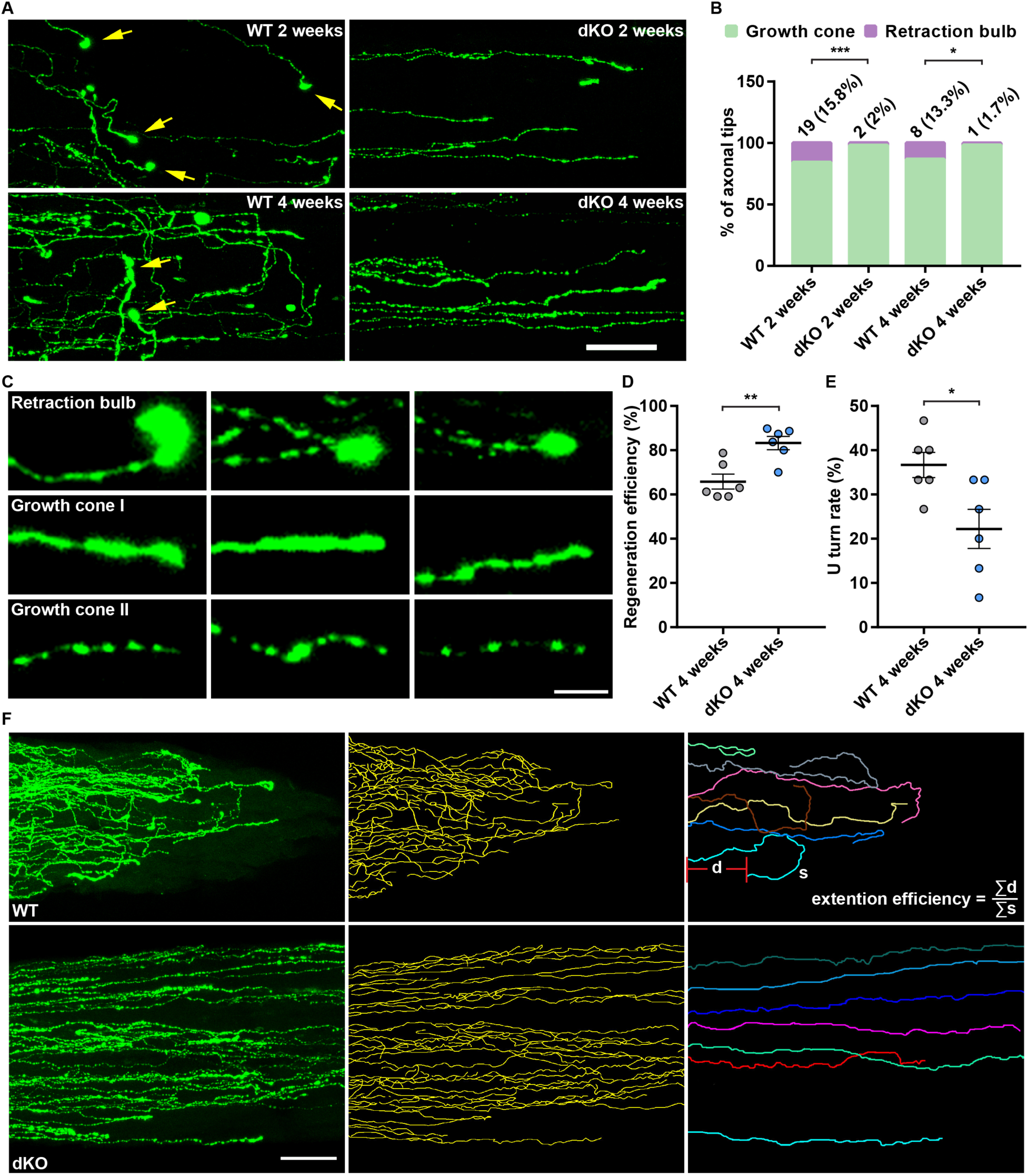
Myosin IIA/B deletion modified cytoskeletal dynamics to promote axon regeneration. (A) Representative images of optic nerves showing that the deletion of myosin IIA/B in RGCs abolished the formation of retraction bulbs in optic nerves 2 and 4 weeks after optic nerve crush. Yellow arrows indicate retraction bulbs. Scale bar, 50 μm. (B) Quantification of retraction bulbs in (A) (Fisher’s exact test, *P* = 0.0004 and 0.0322 for 2 weeks and 4 weeks after optic nerve crush, respectively; n = 120 and 100 axonal tips from 12 nerves in 2-week WT and 10 nerves in dKO groups, respectively, n = 60 axonal tips from 6 nerves in each 4-week group). (C) Representative images of retraction bulbs and growth cones found in different optic nerves. Scale bar, 5 μm. (D) Quantification of axon extension efficiency in (F) and Fig. S5 (unpaired t test, *P* = 0.0032, n = 6 mice in each group, at least 35 axons were analyzed for each mouse). (E) Quantification of U-turn rate in Fig. S6 (unpaired t test, *P* = 0.0210, n = 6 mice in each group, top 15 longest axons were analyzed for each mouse). (F) Left: representative images of optic nerves showing that the deletion of myosin IIA/B in RGCs improved axon extension efficiency 4 weeks after optic nerve crush. Middle: sketches of all axon traces in the left column. Right: detailed trajectories of a few axons (each color represents a single axon) in the left column. As illustrated, the extension efficiency of each nerve was calculated by dividing the summed displacement by the summed length of all traced axons. Scale bar, 50 μm. Data are represented as mean ± SEM. \**P* < 0.05, \*\**P* < 0.01, \*\*\**P* < 0.001. WT, wild type; dKO, double knockout of myosin IIA/B.

The tissue clearing and confocal imaging of whole-mount optic nerves allowed us to visualize the bona fide morphology of regenerating axons. Thus, we next examined how myosin IIA/B knockout influenced the axon extension trajectories 4 weeks after ONC. In wild type nerves, the majority of axons followed a wandering path with many curves and backward turns (U-turns), which resulted in very inefficient axon regeneration towards the distal optic nerve. In contrast, in myosin IIA/B dKO nerves, most regenerating axons were straight with significantly reduced U-turns (Fig. 5D-F and Fig. S5, S6), indicating a higher efficiency of axon regeneration towards the distal end. Together, we think that enhanced optic nerve regeneration induced by myosin IIA/B dKO was achieved through 1) switching retraction bulbs into growth-competent growth cones, and 2) more efficient axon regeneration with straighter axon growth and less U-turns.

### Knocking out non-muscle myosin IIA/B in RGCs after optic nerve crush could also induce optic nerve regeneration

To investigate the translational potentials of myosin IIA/B knockout, we tested if post-injury deletion of myosin IIA/B in RGCs could also promote axon regeneration. We first performed the ONC on wild type and *myosin IIA/B^f/f^* mice, and one day after that, we injected AAV2-Cre into the vitreous humors of these mice. The optic nerve regeneration was assessed 3 weeks after the ONC (Fig. 6A). The result showed that only a small number of axons regenerated for a limited distance in the wild type optic nerves. In contrast, a large number of regenerating axons were observed in dKO optic nerves (Fig. 6B). Most dKO optic nerves had regenerating axons reaching 1500 μm from the crush site, although the most noticeable difference between the two groups was found at 500-1250 μm from the crush site (Fig. 6B). Such a result demonstrated clearly that post-injury treatment with myosin IIA/B knockout could successfully induce axon regeneration in injured optic nerves, indicating myosin IIA/B knockout may potentially be applied to translational practices in treating diseases and injuries involving axon damage.

**Fig. 6.**
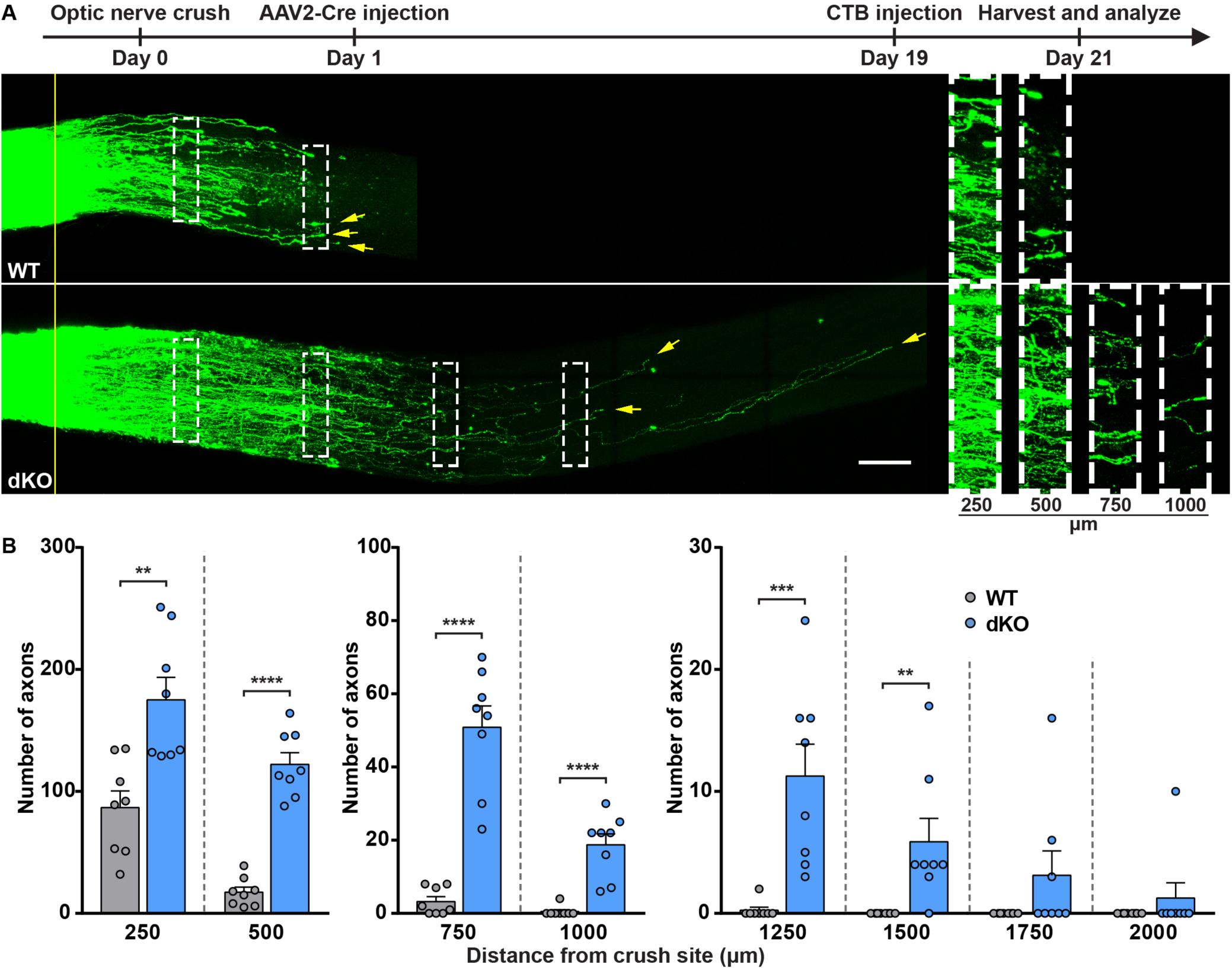
Post-injury deletion of myosin IIA/B could also induce optic nerve regeneration. (A) Top: experimental timeline. Bottom: representative images of optic nerves showing that post-injury deletion of myosin IIA/B in RGCs induced axon regeneration 3 weeks after optic nerve crush. The columns on the right display magnified images of the areas in white dashed boxes on the left, showing axons at 250, 500, 750 and 1000 μm distal to the crush sites. The yellow line indicates the crush sites. Yellow arrows indicate the top 3 longest axons of each nerve. Scale bar, 100 μm (50 μm for the magnified images). (B) Quantification of optic nerve regeneration in (A) (unpaired t test, *P* = 0.0017 at 250 μm, *P* < 0.0001 at 500, 750 and 1000 μm, *P* = 0.0009, 0.0085, 0.1396 and 0.3343 at 1250, 1500, 1750 and 2000 μm, respectively; n = 8 mice in each group).

## Discussion

In addition to diminished intrinsic axon regeneration capacity regulated by changes in gene expression during neuronal maturation, dystrophic growth cone with disruptive cytoskeletal dynamics is another key intrinsic barrier for successful CNS axon regeneration (*18*). Although it has been well recognized that modulation of axonal cytoskeleton would be a plausible approach to enhance CNS axon regeneration, very few studies have shown direct and convincing results. Two previous elegant studies (*34, 35*) have shown that moderate stabilization of microtubules with taxol or epothilone B could promote axon regeneration after the spinal cord injury. The promoting effects were achieved through decreased glial scar formation, which rendered the lesion site more permissive, and improved microtubule protrusion in the growth cones of the injured axons. However, the promoting effects were moderate with regenerating axons only entering the injury site. Similarly, after optic nerve injury low dose taxol treatment alone had little promoting effect on axon regeneration (*36*).

In this study we provided clear evidence that knocking out myosin IIA/B in RGCs alone was sufficient to induce robust and sustained optic nerve regeneration. Our previous in vitro study (*21*) demonstrated that deleting myosin IIA/B acted locally at the growth cone without affecting signaling events at the neuronal soma and gene transcription. Specifically, we showed that inhibiting myosin II activity resulted in reduced level of actin filaments in the growth cone. Retrograde flow of F-actin driven by myosin II acts as a dynamic barrier for microtubule protrusion in the growth cone. As a result, inhibition of myosin II led to significant microtubule protrusion towards the leading edge of the growth cone and increased axon growth rate. Moreover, the promoting effects of myosin II inhibition over inhibitory substrates (CSPGs) occurred within minutes and was reversible, further supporting the effects were local. Indeed, here we carefully examined how RGC axonal morphology and trajectory were affected after myosin IIA/B knockout. The inability of mature CNS axon to form a growth-competent growth cone in the inhibitory environment after injuries is a hallmark of regeneration failure (*18, 20*). In most cases, after CNS injuries, a bulb-like structure is formed at the tip of the injured axon, called the retraction bulb (*33*). In wild type mice, very limited optic nerve regeneration was observed, and retraction bulbs were more often observed at the tips of the axons. In contrast, in myosin IIA/B dKO nerves, very few retraction bulbs were identified, while growth-competent growth cones could be found at almost all axonal tips. The results indicated that deleting myosin IIA/B was an efficient strategy to transform retraction bulbs into growth cones, likely achieved through slowed retrograde flow of actin filaments and the subsequent protrusion of microtubules in the dystrophic growth cones. In addition, confocal microscopy of tissue cleared whole-mount optic nerves allowed us to trace the trajectories of regenerating axons. We found that in the wild type group, the majority of axons showed wandering trajectories with many kinks and U-turns, likely due to the inhibitory substrates in the optic nerve. As a result, such axon extension resulted in very inefficient axon regeneration. When myosin IIA/B were knocked out, most axons followed a straighter path with reduced U-turns, indicating myosin IIA/B deletion could overcome inhibitory cues and greatly enhance regeneration efficiency.

We also examined two well-known signaling pathways in RGCs, mTOR activation and GSK3β inactivation, which occur in the neuronal soma to support the intrinsic axon regeneration ability. The results showed that deleting myosin IIA/B had no effects on these two pathways, suggesting that the intrinsic axon regeneration ability of RGCs might not be elevated. In contrast, overexpression of Lin28a, which promotes optic nerve regeneration by enhancing the intrinsic axon regeneration ability, led to marked activation of mTOR and inactivation of GSK3β in RGCs. Next, by using sensory neuron specific myosin IIA/B conditional knockout mice, we explored how deleting myosin IIA/B affected sensory neuron transcriptome. The results showed that deleting myosin IIA/B did not significantly change the transcription profile in sensory neurons. More importantly, when the transcription levels of many known RAGs were examined, we found that peripheral axotomy significantly changed their mRNA levels, whereas knocking out myosin IIA/B had little effects. Lastly, we directly examined the protein levels of selected RAGs and signaling mediators with western blot. Similarly, knocking out myosin IIA/B had little effects, whereas peripheral axotomy showed significant impact. Together, these results suggested that myosin IIA/B knockout promoted optic nerve regeneration without affecting gene transcription and the intrinsic axon regeneration ability. Collectively, our study suggested that deleting myosin IIA/B promoted optic nerve regeneration by efficiently transforming retraction bulbs into active growth cones and more efficient axon extension within the inhibitory CNS environment.

Long-distance axon regeneration is one of the most important aspects and a prerequisite for successful functional recovery after neural injuries. Several previous studies (*37–39*), including ours (*12*), have shown that combined manipulation of multiple genes/pathways usually had additive or synergistic promoting effects on optic nerve regeneration. Given sufficient time, some axons could even regrow back to their targets in the brain and led to partial function recovery (*39*). Here we showed that combining myosin IIA/B knockout with Lin28a overexpression resulted in surprisingly long-distance optic nerve regeneration. Two weeks after ONC, regenerating axon reached up to 3.25 mm from the crush site (nearly 4 mm in real distance considering 18% shrinkage due to the tissue clearing process (*12*)) without enhancing the RGC survival rate. The longest regenerating axons were about 4.3 mm and close to the optic chiasm. To our knowledge, such a distance was comparable with those in most previously published studies using combinatory approaches. For instance, combined knockout of Pten and SOCS3, together with CNTF, led to optic nerve regeneration up to 3 mm from the crush site 2 weeks after ONC (*37*). Moreover, combination of Zymosan, c-AMP, and Pten deletion could promote optic nerve regeneration to 3 mm from the crush site in 2 weeks (*38*). The effectiveness of the combinatory approach will be optimal if each gene/pathway acts independently. Our previous study found that Lin28 overexpression could induce optic nerve regeneration by enhancing the intrinsic growth ability of RGCs (*12*). A recent study confirmed that specific expression of Lin28 in RGCs could lead to significant optic nerve regeneration. In addition, it also revealed that apart from its direct effect on RGCs, Lin28 specifically expressed in amacrine cells could enhance IGF1-induced optic nerve regeneration by suppressing hyperactivity of amacrine cells induced by optic nerve injury (*13*). As such, Lin28a and myosin II could act in distinct neuronal compartments with diverse cellular mechanisms, resulting in a powerful combination. Because AAV2 also infected amacrine cells, we could not completely rule out the possibility that knocking out myosin IIA/B in amacrine cells somehow contributed to the observed results. Although the number of long-distance regenerating axons observed in this study was relatively low, it was likely due to the poor survival of RGCs, and the different regeneration capacities among RGC subpopulations (*24*). Thus, myosin II knockout can be a new effective option for combination strategies, and future studies combining myosin II knockout with other regeneration approaches, as well as enhanced RGC survival, may potentially lead to large number of axons regenerating back to their original targets in the brain and gain recovery of lost visual function.

Here we also showed that deleting myosin IIA/B in RGCs after the optic nerve injury could induce robust optic nerve regeneration, indicating that myosin IIA/B deletion or inhibition has the potential to be practically used in the treatment of nerve injury. Moreover, the availability of water soluble and stable pharmacological inhibitor of myosin II (*40*) would make future translational applications possible to repair axonal injuries induced by glaucoma, spinal cord injury, traumatic brain injury, and neurodegenerative diseases.

## Materials and Methods

### Mice

All animal experiments were conducted in accordance with the protocol approved by the Institutional Animal Care and Use Committee of the Johns Hopkins University. The *Myh9^f/f^* (stock#032096-UNC) and *Myh10^f/f^* (stock#016981-UNC) mouse strains were obtained from Mutant Mouse Resource and Research Center (MMRRC) at University of North Carolina at Chapel Hill, an NIH-funded strain repository, and were donated to the MMRRC by Robert S. Adelstein, M.D., National Heart, Lung, and Blood Institute (NHLBI). The two lines were crossed to generate *Myh9^f/f^: Myh10^f/f^* mice. *Advillin-Cre* mouse line was a kind gift from Dr. Fan Wang’s laboratory at Duke University, and was crossed with *Myh9^f/f^: Myh10^f/f^* to get *Advillin-Cre: Myh9^f/f^: Myh10^f/f^* conditional knockout mice. The tdTomato reporter line (stock#007909) was purchased from The Jackson Laboratory. Adult mice (6 weeks) of both sexes were used. Genotypes of the mice were determined by PCR using primers provided by MMRRC and The Jackson Laboratory. All animal surgeries were performed under anesthesia induced by intraperitoneal injection of ketamine (100 mg/kg) and xylazine (10 mg/kg) diluted in sterile saline. Details of the surgeries are described below.

### Construct

The pAAV-Ef1a-Lin28a-FLAG plasmid was constructed in a previous study (*12*). Briefly, the Lin28a-FLAG open reading frame with a 5’ BamHI and a 3’ EcoRV restriction sites was synthesized (codon optimized, gBlocks of Integrated DNA Technologies) and used for replacing the EYFP open reading frame in pAAV-Ef1a-EYFP, to obtain the pAAV-Ef1a-Lin28a-FLAG plasmid. pAAV-Ef1a-EYFP was a kind gift from Dr. Hongjun Song’s laboratory at University of Pennsylvania. All restriction enzymes and T4 DNA ligase were purchased from New England Biolabs. Plasmids were amplified using DH5α competent cells (Thermo Fisher Scientific) and purified with Endofree plasmid maxi kit (Qiagen).

### Optic nerve regeneration model

Intravitreal viral injection, optic nerve crush and RGC axon labeling were performed as previously described (*9*). Briefly, under anesthesia, 1.5 μl of AAV2 virus was injected into the right vitreous humor of a mouse with a Hamilton syringe (32-gauge needle). The position and direction of the injection were well-controlled to avoid injury to the lens. Two weeks later, the right optic nerve of the mouse was exposed intraorbitally and crushed with Dumont #5 fine forceps (Fine Science Tools) for 5 s at approximately 1 mm behind the optic disc. To label RGC axons in the optic nerve, 1.5 μl of Alexa Fluor 594-conjugated CTB (2 μg/μl, Thermo Fisher Scientific) was injected into the right vitreous humor with a Hamilton syringe (32-gauge needle) 2 days before the mouse was sacrificed by transcardial perfusion under anesthesia. The right optic nerve and bilateral retinas were dissected out and post-fixed in 4% PFA overnight at 4°C. AAV2-Cre (SL100813) was purchased from SignaGen Laboratories. AAV2-Lin28a-FLAG was also packaged by SignaGen Laboratories. All viruses used had titers over 1 x 10^13^ gc/ml.

For post-injury treatment model, all procedures were done in the same way except the intravitreal viral injection was conducted one day after the optic nerve crush.

### Optic nerve dehydration and clearing

Dehydration and clearing of optic nerves were done based on previous studies (*14, 41*). Briefly, fixed optic nerves were first dehydrated in increasing concentrations of tetrahydrofuran (50%, 70%, 80%, 100% and 100%, v/v % in distilled water, 20 min each, Sigma-Aldrich) and then cleared in a solution of benzyl alcohol and benzyl benzoate (BABB, 1:2, Sigma-Aldrich). Incubations were done on an orbital shaker at room temperature. The nerves were stored in BABB in the dark at room temperature.

### Analysis of RGC axon regeneration

Tissue cleared whole-mount optic nerves were imaged with a 20x objective on a Zeiss 800 confocal microscope. For each optic nerve, Z-stack and tiling (10% overlap) functions were used to acquire stacked 2-μm-thick planes of the whole area of interest and the tiles were stitched.

To quantify the number of regenerating axons in each optic nerve, every 8 consecutive planes were Z-projected (maximum intensity) to generate a series of Z-projection images of 16-μm-thick optical sections. At each 250-μm interval from the crush site, the number of CTB-labeled axons was counted in each Z-projection image and summed over all optical sections.

To quantify the average length of top 5 longest axons of each optic nerve, all stitched 2-μm-thick planes were Z-projected (maximum intensity) to obtain a single Z-projection image of the nerve. Top 5 longest regenerating axons were manually traced from the axonal tips to the crush site using the Fiji software (NIH) to acquire the lengths of the axons.

### Immunohistochemistry of whole-mount retinas

Fixed retinas were first radially cut into a petal shape (4 incisions) and blocked with PBST (1%) containing 10% goat serum for 1 hr at room temperature. The retinas were then sequentially stained with primary antibodies overnight at 4°C, and corresponding Alexa Fluor-conjugated secondary antibodies (1:500, Thermo Fisher Scientific) for 2 hr at room temperature. All antibodies were diluted with the blocking buffer. Following each antibody incubation, the retinas were washed with PBST (0.3%) for 4 times (15 min each). After the last wash, the retinas were mounted onto slides with Fluoroshield (Sigma-Aldrich). Fluorescent images of the flat-mounted retinas were acquired with a 20x objective on a Zeiss 800 confocal microscope.

### Analysis of RGC transduction rate

To quantify RGC transduction rate, uninjured right retinas (no optic nerve crush) were taken from transcardially perfused *Myh9^f/f^: Myh10^f/f^* mice 2 weeks after intravitreal AAV2-Cre injection. The retinas were stained with mouse anti-tubulin β3 (Tuj1, 1:500, BioLegend) and rabbit anti-Cre recombinase (1:100, Cell Signaling Technology) antibodies following the steps mentioned above (see Immnunohistochemistry of whole-mount retinas). Five to eight fields under 20x objective were randomly obtained from the peripheral regions of each flat-mounted retina. For each mouse, RGC transduction rate was calculated by dividing the total number of Cre^+^/Tuj1^+^ cells in all fields by the total number of Tuj1^+^ cells in all fields. Only cells in the ganglion cell layer were counted.

### Analysis of RGC survival rate

To quantify RGC survival rate, C57Bl6/J and *Myh9^f/f^: Myh10^f/f^* mice injected with AAV2-Cre were transcardially perfused 2 weeks after optic nerve crush and both retinas of each mouse were collected. The retinas were stained with mouse anti-tubulin β3 antibody (Tuj1, 1:500, BioLegend) following the steps mentioned above (see Immnunohistochemistry of whole-mount retinas). Seven or eight fields under 20x objective were randomly taken from the peripheral regions of each flat-mounted retina. For each mouse, RGC survival rate was calculated by dividing the average number of Tuj1^+^ cells in one field in the injured retina (right) by that in the uninjured retina (left). Only cells in the ganglion cell layer were counted.

### Immunohistochemistry of retinal sections

Fixed retinas were sectioned with a cryostat (10 μm) and the retinal sections were warmed on a slide warmer at 37°C for 1 hr. Sections were rinsed once in PBS, soaked in 100°C citrate buffer (pH 6) for 15 min, let to cool in the buffer to room temperature and then washed twice (5 min each) in PBS. After being blocked with PBST (0.3%) containing 10% goat serum at room temperature for 1 hr, the sections were stained with primary antibodies against target molecules overnight at 4 °C, followed by corresponding Alexa Fluor-conjugated secondary antibodies (1:500, Thermo Fisher Scientific) at room temperature for 1 hr. All antibodies were diluted with the blocking buffer. The sections were washed for 4 times (5, 5, 10, 10 min) with PBST (0.3%) following each antibody incubation and finally mounted with DAPI Fluoromount-G (SouthernBiotech). Fluorescent images of the retinal sections were taken with a CCD camera connected to a Zeiss inverted fluorescence microscope controlled by AxioVision software.

### Analysis of myosin IIB level in RGCs

To analyze myosin IIB level in RGCs, both retinas (uninjured) of each mouse were taken from transcardially perfused *Myh9^f/f^: Myh10^f/f^* mice 2 weeks after intravitreal AAV2-Cre injection and sectioned. The retinal sections were stained with mouse anti-tubulin β3 (Tuj1, 1:500, BioLegend) and rabbit anti-myosin IIB (1:100, Thermo Fisher Scientific) antibodies following the steps mentioned above (see Immnunohistochemistry of retinal sections).

To quantify the fluorescence intensity of myosin IIB in all RGCs, at least 7 non-adjacent retinal sections acquired with identical imaging configurations were analyzed for each retina. Fluorescence intensity was measured using the “outline spline” function of AxioVision and the background fluorescence intensity was subtracted.

### Analysis of S6 and GSK3β phosphorylation

To analyze S6 and GSK3β phosphorylation in RGCs, C57Bl6/J and *Myh9^f/f^: Myh10^f/f^* mice injected with AAV2-Cre, and *Myh9^f/f^: Myh10^f/f^* mice injected with AAV2-Lin28a-FLAG were transcardially perfused 2 weeks after optic nerve crush and the right retina of each mouse was collected and sectioned. The retinal sections were stained with mouse anti-tubulin β3 antibody (Tuj1, 1:500, BioLegend), and rabbit anti-pS6 Ser235/236 (1:200, Cell Signaling Technology) or rabbit anti-pGSK3β Ser9 (1:200, Cell Signaling Technology) antibody following the steps mentioned above (see Immnunohistochemistry of retinal sections).

To quantify the percentage of pS6^+^ or pGSK3β^+^ RGCs, at least 363 or 434 RGCs from at least 7 non-adjacent retinal sections from each mouse were analyzed. For each mouse, the percentage of pS6^+^ or pGSK3β^+^ RGCs was calculated by dividing the number of pS6^+^/Tuj1^+^ or pGSK3β^+^/Tuj1^+^ cells by the number of Tuj1^+^ cells. Only cells in the ganglion cell layer were counted.

The relative fluorescence intensity of each RGC was calculated by dividing the fluorescence intensity of the RGC by that of its adjacent tissue. To quantify the fluorescence intensity of pS6 or pGSK3β in all RGCs, 20 retinal sections acquired with identical imaging configurations from at least 2 mice were analyzed for each group. To quantify the fluorescence intensity of pS6 or pGSK3β in S6-activated or GSK3β-inactivated RGCs, at least 40 RGCs with identical imaging configurations from at least 2 mice were analyzed for each group. Fluorescence intensity of RGCs were measured using the “outline spline” function of AxioVision.

### Sciatic nerve injury model

Under anesthesia, bilateral sciatic nerves of a mouse were exposed and transected with spring scissors right below pelvis. Nerves were only exposed but not transected for a sham surgery. Three days after the surgery, the mouse was euthanized and bilateral L4/5 DRGs were collected and used for total RNA or protein extraction.

### mRNA sequencing and data analysis

Total RNA was isolated with RNeasy mini kit (Qiagen) and RNA integrity was determined by Agilent 2100 bioanalyzer and RNA 6000 Nano kit (Agilent Technologies). Paired-end libraries were synthesized using the TruSeq RNA library preparation kit (Illumina). Briefly, mRNA molecules were purified using oligo dT-attached magnetic beads. Following purification, the mRNA molecules were fragmented into small pieces using divalent cations at 94°C for 8 min. The cleaved RNA fragments were reversely transcribed into first strand cDNA using reverse transcriptase and random primers. This was followed by second strand cDNA synthesis using DNA Polymerase I and RNase H. These cDNA fragments then went through an end repair process, the addition of a single base, and the ligation of the adapters. The products were then purified and enriched with PCR to obtain the final cDNA library. Purified libraries were quantified by Qubit 2.0 fluorometer (Thermo Fisher Scientific) and validated by Agilent 2100 bioanalyzer (Agilent Technologies) to confirm the insert size and calculate the mole concentration. Cluster was generated by cBot with the library diluted to 10 pM and sequencing was done on HiSeq 2500 (Illumina).

Sequencing raw reads were preprocessed by filtering out rRNA reads, sequencing adapters, short-fragment reads and other low-quality reads using Seqtk (github.com/lh3/seqtk). Hisat2 (version 2.0.4) (*42*) was used to map the cleaned reads to the mouse GRCm38.p4 (mm10) reference genome with two mismatches. After genome mapping, Stringtie (version 1.3.0) (*43, 44*) was run with a reference annotation to generate FPKM (Fragments Per Kilobase of exon model per Million mapped reads) values for known gene models (*45*). Differentially expressed genes were identified using edgeR (*46*). The *P* value significance threshold in multiple tests was set by the false discovery rate (FDR) (*47, 48*). The cut-off for differentially expressed genes was set as *P* < 0.05 and |log2fold change| > 1. For genes used for gene ontology (GO) analyses, an additional cut-off of average FPKM > 1 in at least one condition was applied. GO analysis was done using DAVID Bioinformatics Resources 6.8 (*49, 50*). GO terms came from biological process, cellular component and molecular function categories.

### Western blot analysis

Total protein was extracted from L4/5 DRGs or retinas using the RIPA buffer containing protease inhibitor cocktail (Sigma-Aldrich) and phosphatase inhibitor cocktail (Sigma-Aldrich). Identical amount of total protein from each condition was then separated by 4-12% gradient SDS-PAGE gel electrophoresis and transferred onto polyvinylidene fluoride membranes. After being blocked with TBST (1%) containing 5% blotting-grade blocker (Bio-Rad), the membranes were incubated overnight with primary antibodies against target molecules at 4 °C, followed by corresponding HRP-linked secondary antibodies (1:2000, Cell Signaling Technology) for 1 hr at room temperature. All antibodies were diluted with blocking buffer. The membranes were washed with TBST (1%) for four times (5, 5, 10, 10 min) after each antibody incubation. Rabbit primary antibodies against myosin IIA (1:1000), Atf3 (1:1000), Gap43 (1:1000), c-Jun (1:1000), c-Myc (1:1000), pAkt Ser473 (1:2000), pGSK3β Ser9 (1:1000), pS6 Ser235/236 (1:2000) were purchased from Cell Signaling Technology. Mouse anti-β-actin primary antibody (1:5000) was from Sigma-Aldrich.

### Analysis of axonal tip morphology

The method was derived from a previous study (*33*). For each optic nerve, top 10 longest axons were first identified in the Z-projection image of the nerve. Then the maximum diameter of each axonal tip and the diameter of the cylindrical shaft of the corresponding axon were measured using Fiji software (NIH), and a tip/shaft ratio was calculated. An axonal tip was defined as a retraction bulb if its tip/shaft ratio was over 4. Otherwise, it was defined as a growth cone.

### Analysis of axon extension efficiency

For each optic nerve, a 250-μm-long region with equivalent number of axons in the Z-projection image of the nerve was used for analysis. All traceable axons within this region were manually traced. For each axon, the length (covered distance on its trajectory) and the displacement (distance along the longitudinal axis of the nerve, sometimes could be zero or negative) between the start point and the end point were measured using Fiji software (NIH). The extension efficiency of each nerve was calculated by dividing the summed displacement by the summed length of all axons.

### Analysis of U-turn rate

For each optic nerve, top 15 longest axons were identified in the Z-projection image of the nerve and their trajectories near the axonal tips were traced. A U-turn was defined when the angle between the final direction of the axonal tip and the positive longitudinal axis of the optic nerve was larger than 90 degrees. U-turn rate of each nerve was calculated by dividing the number of axons that made U-turns by 15.

### Quantification and statistical analysis

Statistical analyses were done with GraphPad Prism 7 and the significance level was set as *P* < 0.05. Data are represented as mean ± SEM unless specifically stated. For comparisons between two groups, two-tailed unpaired or paired t test was used. For comparisons among three or more groups, one-way ANOVA followed by Tukey’s multiple comparisons test was used to determine the statistical significance. Fisher’s exact test was used to test contingency tables. All details regarding statistical analyses, including the tests used, *P* values, exact values of n, definitions of n, are described in figure legends.

## Acknowledgements

We appreciate Dr. Michele Pucak’s help in confocal imaging experiments and analyses.

## Funding

The study was supported by grants (to F.Q.Z.) from NIH (R01NS064288, R01NS085176, R01GM111514, R01EY027347), the Craig H. Neilsen Foundation (259450), and the BrightFocus Foundation (G2017037). Saijilafu was supported by the grants from the National Natural Science Foundation of China (81571189, 81772353), the National Key Research and Development Program (2016YFC1100203), and Innovation and Entrepreneurship Program of Jiangsu Province.

## Author contributions

X.-W.W., S.-G.Y., S., and F.-Q.Z. conceived the study and designed the project; X.-W.W. and S.-G.Y. performed most of the experiments; C.Z. performed and analyzed the immunostaining experiments; Y.Z. analyzed optic nerve regeneration; C.Z. and Y.Z. analyzed the axon morphologies and trajectories; J.-J.M. performed the sensory neuron RNA-seq experiment; Y.-L.W. and G.-L.M. helped with the optic nerve regeneration experiments; B.-B.Y., and A.R.K. helped with the data analyses; X.-W.W. and F.-Q.Z. wrote the manuscript with contribution from all authors.

## Competing interests

The authors declare no competing interests.

## Data and material availability

The data that support the findings of this study are available from the authors upon request. RNA-seq raw data will be deposited and accession code will be available before publication.

---

**Fig. S1.**
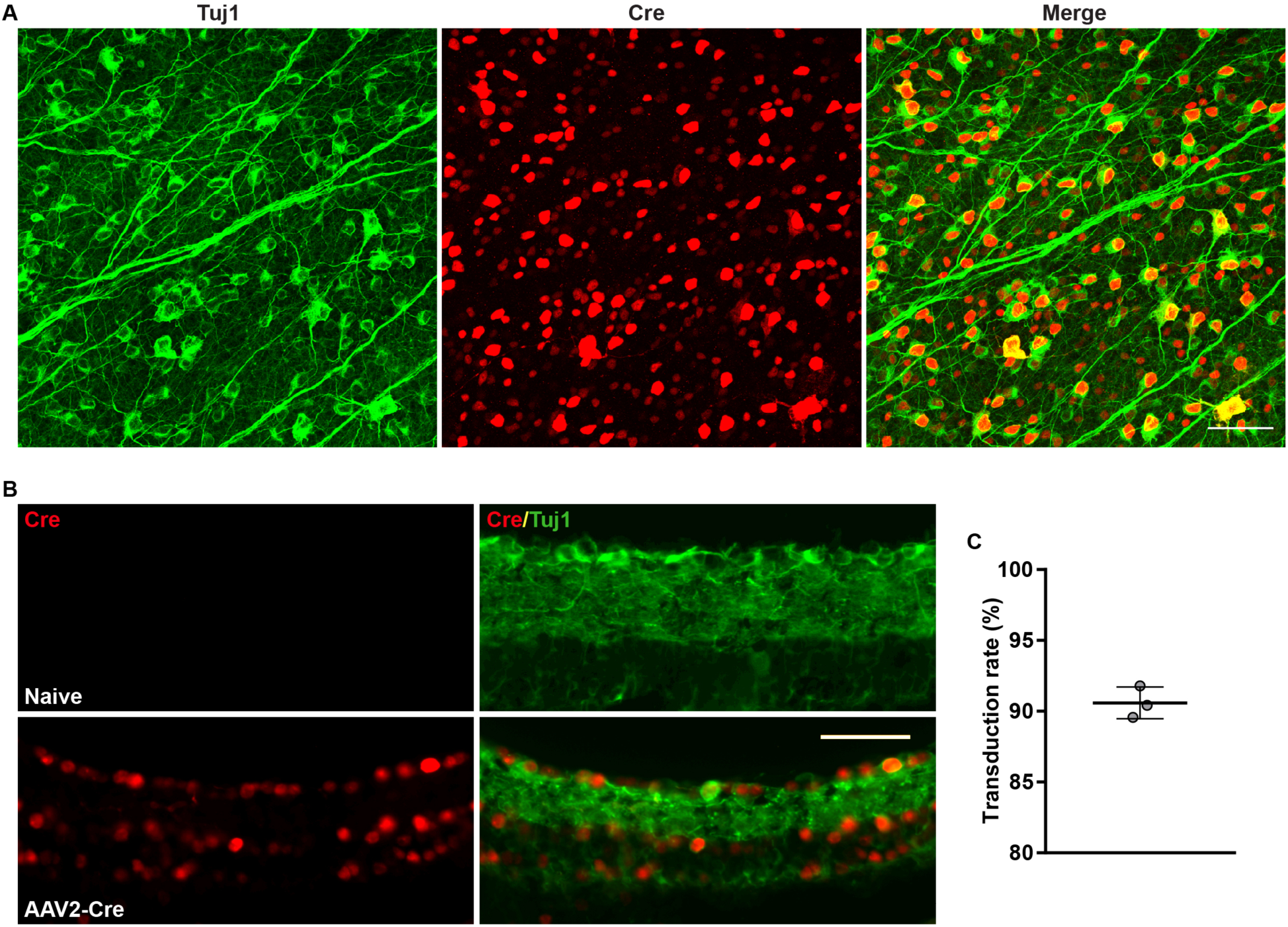
Transduction rate of AAV2-Cre. Related to Fig. 1. (A) Representative images of flat-mounted retinas showing the high transduction rate of AAV2-Cre in RGCs. Flat-mounted retinas were stained with anti-tubulin β3 (Tuj1, green) and anti-Cre recombinase (red) antibodies. Scale bar, 50 μm. (B) Representative images of retinal sections verifying the specificity of the anti-Cre antibody and the high transduction rate of AAV2-Cre in RGCs. Retinal sections were stained with anti-tubulin β3 (Tuj1, green) and anti-Cre (red) antibodies. Scale bar, 50 μm. (C) Quantification of the transduction rate of AAV2-Cre in RGCs in (A). The average transduction rate was 90.59 ± 1.114% (n = 3 mice, 5-8 fields were analyzed for each mouse, data are represented as mean ± SD).

**Fig. S2.**
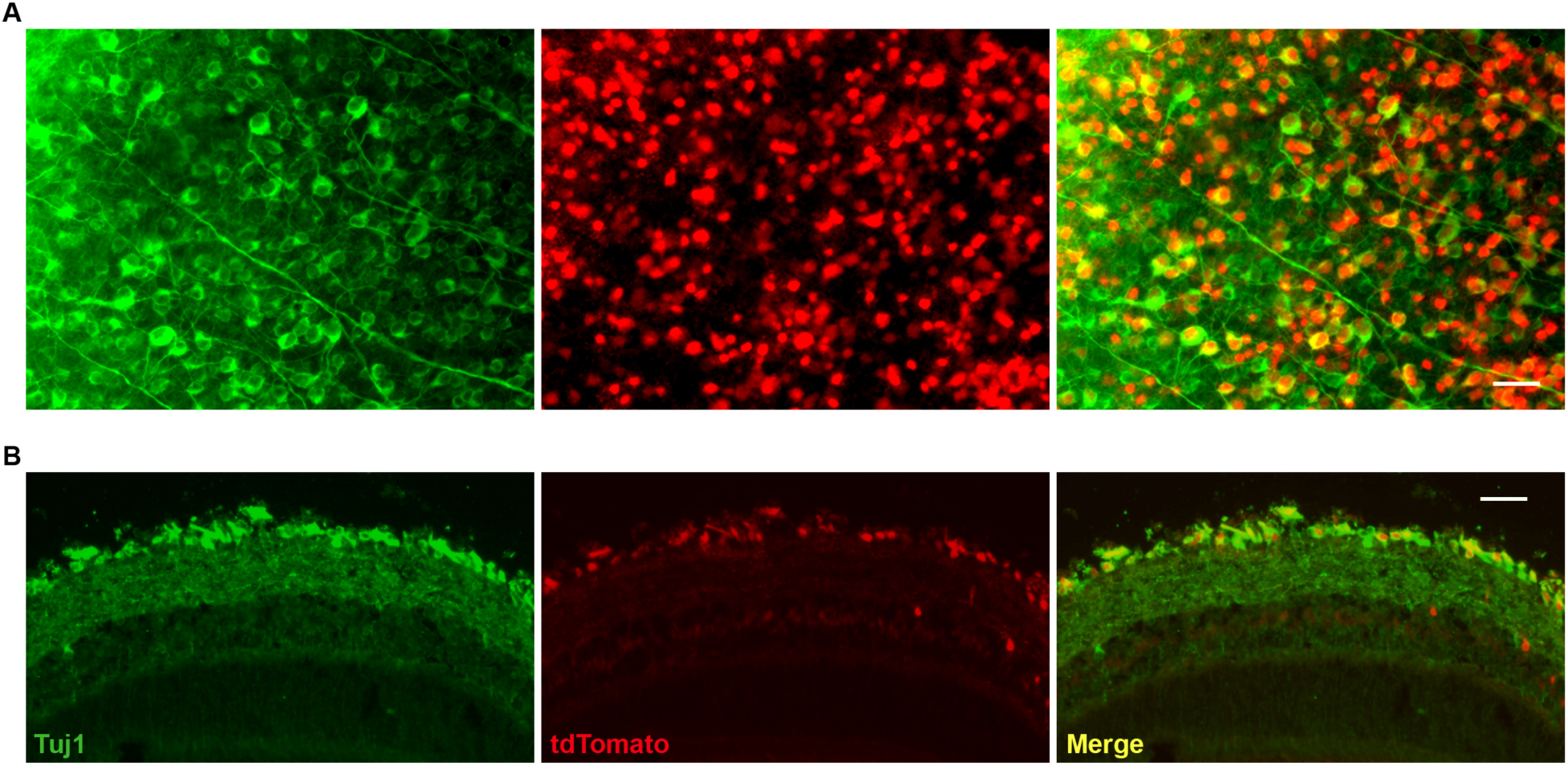
Verification of the Cre-mediated gene recombination. Related to Fig. 1. (A, B) Representative images of flat-mounted retinas (A) and retinal sections (B) showing the expression of tdTomato in RGCs of tdTomato reporter mice, indicating successful Cre-mediated gene recombination 2 weeks after intravitreal AAV2-Cre injection. Flat-mounted retinas and retinal sections were stained with anti-tubulin β3 (Tuj1, green). Scale bar, 50 μm.

**Fig. S3.**
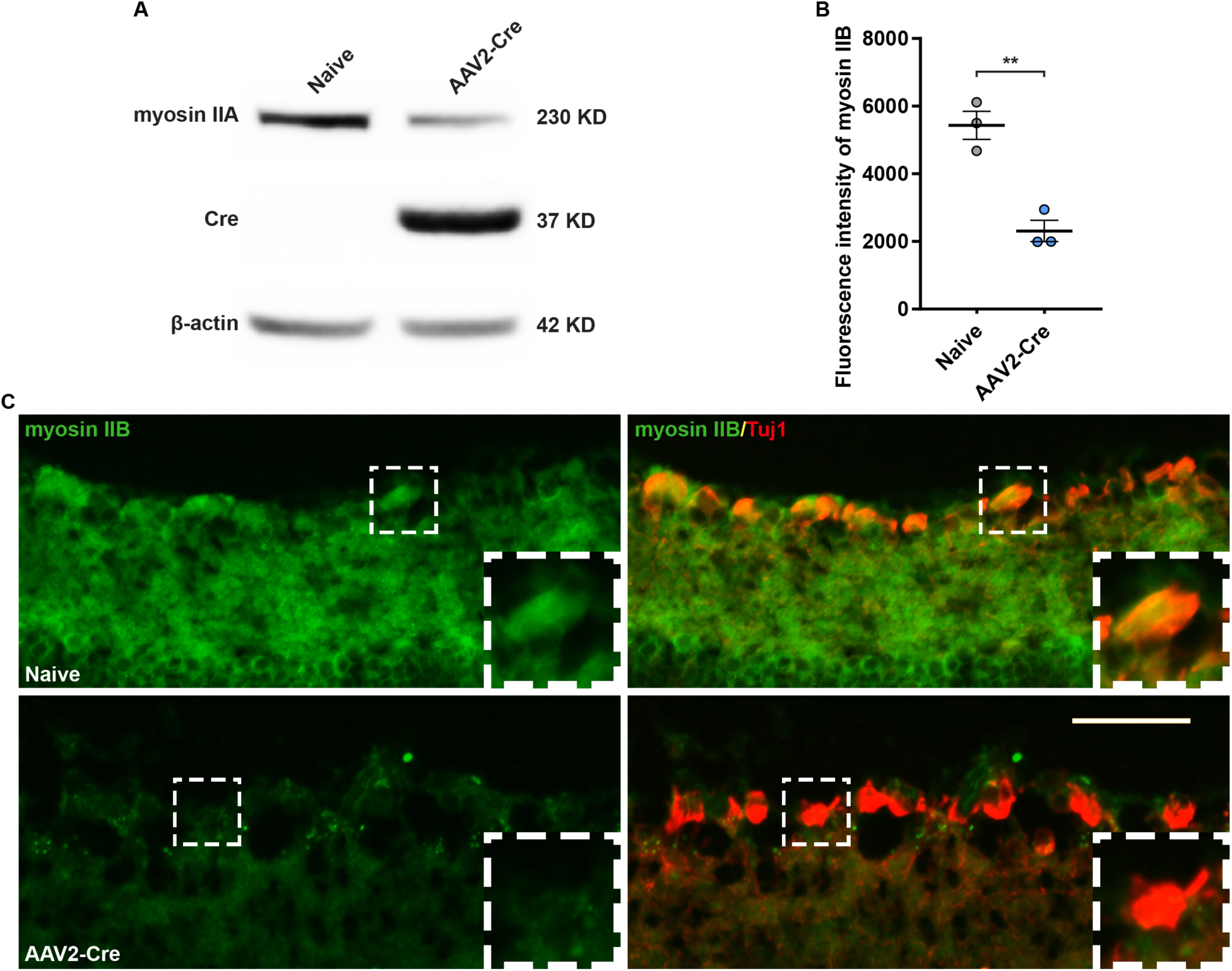
Verification of myosin IIA/B deletion. Related to Fig. 1. (A) Western blot result showing the decreased protein level of myosin IIA in retina of a *myosin IIA/B^f/f^* mouse 2 weeks after intravitreal AAV2-Cre injection. (B) Quantification of average fluorescence intensity of myosin IIB in (C) (paired t test, *P* = 0.0057, n = 3 mice, data are represented as mean ± SEM, \*\**P* < 0.01, all images were taken with identical configurations). (C) Representative images of retinal sections from the naïve (top) or the AAV2-Cre injected retina (bottom) of a *myosin IIA/B^f/f^* mouse showing the successful deletion of myosin IIB in RGCs 2 weeks after intravitreal AAV2-Cre injection. The insets display magnified images of the RGCs marked in white dashed boxes. Retinal sections were stained with anti-tubulin β3 (Tuj1, red) and anti-myosin IIB (green) antibodies. Scale bar, 50 μm (25 μm for the magnified images).

**Fig. S4.**
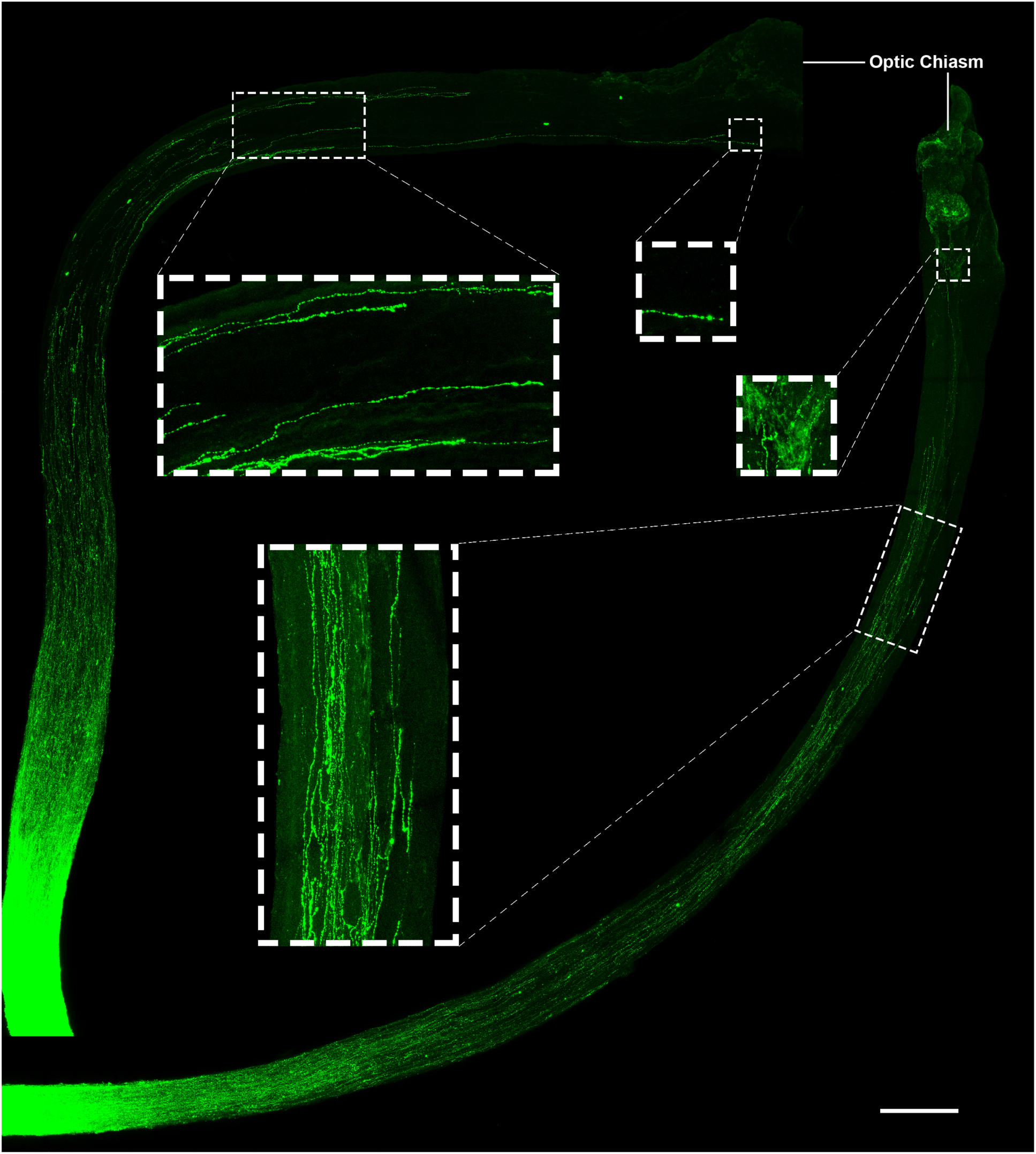
Combination of myosin IIA/B deletion and Lin28a overexpression greatly enhanced optic nerve regeneration up to the optic chiasm. Related to Fig. 2. Images of two optic nerves in the combinatory treatment group with longest axons reaching optic chiasm. Magnified images show detailed morphologies of axons in white dashed boxes. Scale bar, 200 μm (66.7 μm for the magnified images).

**Fig. S5.**
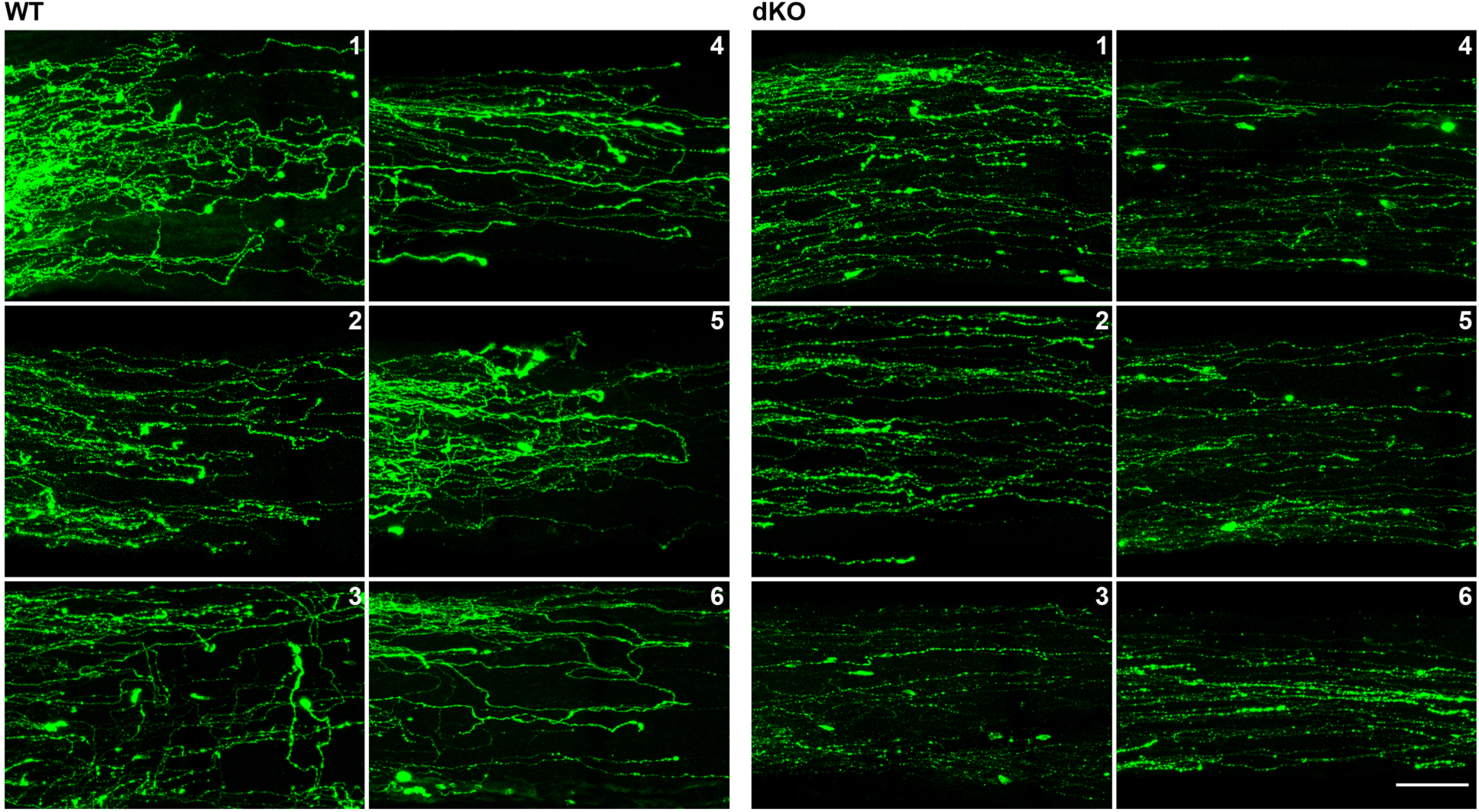
Deletion of myosin IIA/B in RGCs improved axon extension efficiency. Related to Fig. 5. Images of the regions used for the quantification of axon extension efficiency (one region from each optic nerve). Scale bar, 50 μm. WT, wild type; dKO, double knockout of myosin IIA/B.

**Fig. S6.**
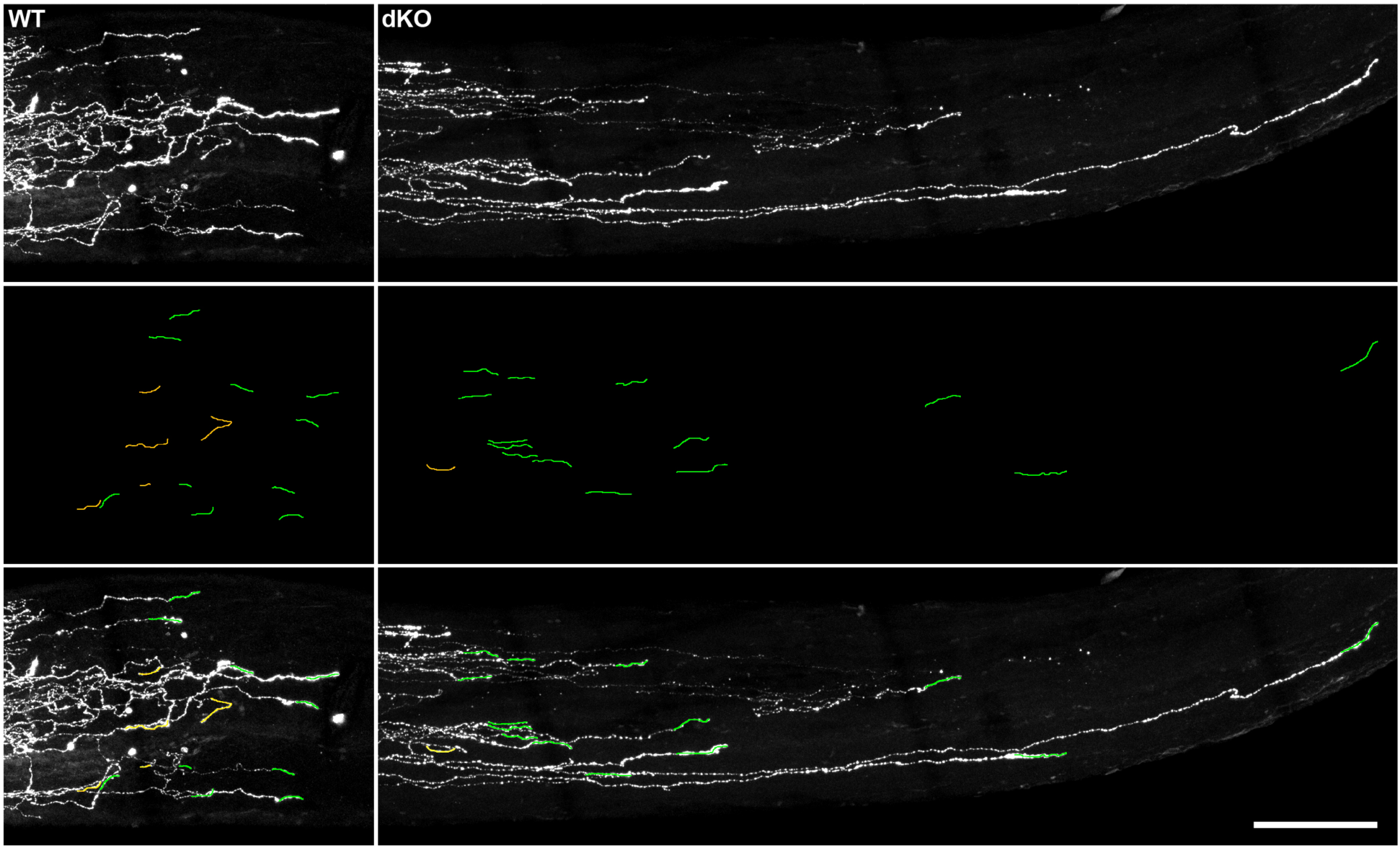
Deletion of myosin IIA/B reduced U-turn rate. Related to Fig. 5. Top: representative images of the leading-edge area of optic nerves showing that myosin IIA/B deletion reduced U-turn rate. Middle: axon trajectories near axonal tips, U-turns are labeled in yellow. Bottom: overlay of top and middle rows. Scale bar, 50 μm. WT, wild type; dKO, double knockout of myosin IIA/B.

**Movie S1. A 3D animation of regenerating axons in an optic nerve.**

## Notes

#### Summary of Updates

We have added the new results that knocking out myosin IIA/B in RGCs after optic nerve crush also led to significant optic nerve regeneration. The results provided proof-of-concept evidence regarding potential translational application of the findings.

## Reference and Notes

1. Z. He, Y. Jin, Intrinsic Control of Axon Regeneration. Neuron 90, 437–451 (2016).

2. M. Curcio, F. Bradke, Axon Regeneration in the Central Nervous System: Facing the Challenges from the Inside. Annu Rev Cell Dev Biol 34, 495–521 (2018).

3. S. David, A. J. Aguayo, Axonal elongation into peripheral nervous system “bridges” after central nervous system injury in adult rats. Science 214, 931–933 (1981).

4. J. W. Fawcett, The Paper that Restarted Modern Central Nervous System Axon Regeneration Research. Trends Neurosci 41, 239–242 (2018).

5. P. M. Richardson, U. M. Mcguinness, A. J. Aguayo, Axons from Cns Neurons Regenerate into Pns Grafts. Nature 284, 264–265 (1980).

6. K. F. So, A. J. Aguayo, Lengthy regrowth of cut axons from ganglion cells after peripheral nerve transplantation into the retina of adult rats. Brain Res 328, 349–354 (1985).

7. C. G. Geoffroy, B. Zheng, Myelin-associated inhibitors in axonal growth after CNS injury. Current opinion in neurobiology 27, 31–38 (2014).

8. J. K. Lee, B. Zheng, Role of myelin-associated inhibitors in axonal repair after spinal cord injury. Exp Neurol 235, 33–42 (2012).

9. K. K. Park et al., Promoting axon regeneration in the adult CNS by modulation of the PTEN/mTOR pathway. Science 322, 963–966 (2008).

10. P. D. Smith et al., SOCS3 Deletion Promotes Optic Nerve Regeneration In Vivo. Neuron 64, 617–623 (2009).

11. D. L. Moore et al., KLF family members regulate intrinsic axon regeneration ability. Science 326, 298–301 (2009).

12. X. W. Wang et al., Lin28 Signaling Supports Mammalian PNS and CNS Axon Regeneration. Cell Rep 24, 2540–2552 e2546 (2018).

13. Y. Zhang et al., Elevating Growth Factor Responsiveness and Axon Regeneration by Modulating Presynaptic Inputs. Neuron 103, 39–51 e35 (2019).

14. X. Luo et al., Three-dimensional evaluation of retinal ganglion cell axon regeneration and pathfinding in whole mouse tissue after injury. Exp Neurol 247, 653–662 (2013).

15. V. Pernet et al., Misguidance and modulation of axonal regeneration by Stat3 and Rho/ROCK signaling in the transparent optic nerve. Cell Death Dis 4, e734 (2013).

16. K. Liu et al., PTEN deletion enhances the regenerative ability of adult corticospinal neurons. Nat Neurosci 13, 1075–1081 (2010).

17. C. G. Geoffroy, B. J. Hilton, W. Tetzlaff, B. Zheng, Evidence for an Age-Dependent Decline in Axon Regeneration in the Adult Mammalian Central Nervous System. Cell Rep 15, 238–246 (2016).

18. O. Blanquie, F. Bradke, Cytoskeleton dynamics in axon regeneration. Current opinion in neurobiology 51, 60–69 (2018).

19. E. M. Hur et al., GSK3 controls axon growth via CLASP-mediated regulation of growth cone microtubules. Genes Dev 25, 1968–1981 (2011).

20. E. M. Hur, Saijilafu, F. Q. Zhou, Growing the growth cone: remodeling the cytoskeleton to promote axon regeneration. Trends in Neurosciences 35, 164–174 (2012).

21. E. M. Hur et al., Engineering neuronal growth cones to promote axon regeneration over inhibitory molecules. Proc Natl Acad Sci U S A 108, 5057–5062 (2011).

22. X. Guo, W. D. Snider, B. Chen, GSK3beta regulates AKT-induced central nervous system axon regeneration via an eIF2Bepsilon-dependent, mTORC1-independent pathway. Elife 5, e11903 (2016).

23. L. Miao et al., mTORC1 is necessary but mTORC2 and GSK3beta are inhibitory for AKT3-induced axon regeneration in the central nervous system. Elife 5, (2016).

24. X. Duan et al., Subtype-specific regeneration of retinal ganglion cells following axotomy: effects of osteopontin and mTOR signaling. Neuron 85, 1244–1256 (2015).

25. S. Li et al., Promoting axon regeneration in the adult CNS by modulation of the melanopsin/GPCR signaling. Proc Natl Acad Sci U S A 113, 1937–1942 (2016).

26. W. Pita-Thomas, M. Mahar, A. Joshi, D. Gan, V. Cavalli, HDAC5 promotes optic nerve regeneration by activating the mTOR pathway. Exp Neurol 317, 271–283 (2019).

27. J. H. Lim et al., Neural activity promotes long-distance, target-specific regeneration of adult retinal axons. Nat Neurosci 19, 1073–1084 (2016).

28. H. Tsujino et al., Activating transcription factor 3 (ATF3) induction by axotomy in sensory and motoneurons: A novel neuronal marker of nerve injury. Mol Cell Neurosci 15, 170–182 (2000).

29. K. Tanabe, I. Bonilla, J. A. Winkles, S. M. Strittmatter, Fibroblast growth factor-inducible-14 is induced in axotomized neurons and promotes neurite outgrowth. Journal of Neuroscience 23, 9675–9686 (2003).

30. K. J. Christie, C. A. Webber, J. A. Martinez, B. Singh, D. W. Zochodne, PTEN Inhibition to Facilitate Intrinsic Regenerative Outgrowth of Adult Peripheral Axons. Journal of Neuroscience 30, 9306–9315 (2010).

31. A. Apara et al., KLF9 and JNK3 Interact to Suppress Axon Regeneration in the Adult CNS. Journal of Neuroscience 37, 9632–9644 (2017).

32. Y. Sekine et al., Functional Genome-wide Screen Identifies Pathways Restricting Central Nervous System Axonal Regeneration. Cell Reports 23, 415–428 (2018).

33. A. Erturk, F. Hellal, J. Enes, F. Bradke, Disorganized microtubules underlie the formation of retraction bulbs and the failure of axonal regeneration. J Neurosci 27, 9169–9180 (2007).

34. F. Hellal et al., Microtubule Stabilization Reduces Scarring and Causes Axon Regeneration After Spinal Cord Injury. Science 331, 928–931 (2011).

35. J. Ruschel, et al., Axonal regeneration. Systemic administration of epothilone B promotes axon regeneration after spinal cord injury. Science 348, 347–352 (2015).

36. V. Sengottuvel, M. Leibinger, M. Pfreimer, A. Andreadaki, D. Fischer, Taxol facilitates axon regeneration in the mature CNS. J Neurosci 31, 2688–2699 (2011).

37. F. Sun et al., Sustained axon regeneration induced by co-deletion of PTEN and SOCS3. Nature 480, 372–375 (2011).

38. T. Kurimoto et al., Long-distance axon regeneration in the mature optic nerve: contributions of oncomodulin, cAMP, and pten gene deletion. The Journal of neuroscience: the official journal of the Society for Neuroscience 30, 15654–15663 (2010).

39. S. de Lima et al., Full-length axon regeneration in the adult mouse optic nerve and partial recovery of simple visual behaviors. Proc Natl Acad Sci U S A 109, 9149–9154 (2012).

40. B. H. Varkuti et al., A highly soluble, non-phototoxic, non-fluorescent blebbistatin derivative. Sci Rep 6, 26141 (2016).

41. A. Erturk et al., Three-dimensional imaging of solvent-cleared organs using 3DISCO. Nat Protoc 7, 1983–1995 (2012).

42. D. Kim, B. Langmead, S. L. Salzberg, HISAT: a fast spliced aligner with low memory requirements. Nat Methods 12, 357–360 (2015).

43. M. Pertea et al., StringTie enables improved reconstruction of a transcriptome from RNA-seq reads. Nat Biotechnol 33, 290–295 (2015).

44. M. Pertea, D. Kim, G. M. Pertea, J. T. Leek, S. L. Salzberg, Transcript-level expression analysis of RNA-seq experiments with HISAT, StringTie and Ballgown. Nat Protoc 11, 1650–1667 (2016).

45. A. Mortazavi, B. A. Williams, K. McCue, L. Schaeffer, B. Wold, Mapping and quantifying mammalian transcriptomes by RNA-Seq. Nat Methods 5, 621–628 (2008).

46. M. D. Robinson, D. J. McCarthy, G. K. Smyth, edgeR: a Bioconductor package for differential expression analysis of digital gene expression data. Bioinformatics 26, 139–140 (2010).

47. Y. Benjamini, Y. Hochberg, Controlling the False Discovery Rate - a Practical and Powerful Approach to Multiple Testing. J R Stat Soc B 57, 289–300 (1995).

48. Y. Benjamini, D. Yekutieli, The control of the false discovery rate in multiple testing under dependency. Ann Stat 29, 1165–1188 (2001).

49. D. W. Huang, B. T. Sherman, R. A. Lempicki, Systematic and integrative analysis of large gene lists using DAVID bioinformatics resources. Nature Protocols 4, 44–57 (2009).

50. D. W. Huang, B. T. Sherman, R. A. Lempicki, Bioinformatics enrichment tools: paths toward the comprehensive functional analysis of large gene lists. Nucleic Acids Research 37, 1–13 (2009).

